# Placental gene expression-based cell type deconvolution: Cell proportions drive preeclampsia gene expression differences

**DOI:** 10.1101/2021.07.29.454041

**Authors:** Kyle A Campbell, Justin A Colacino, Muraly Puttabyatappa, John F Dou, Elana R Elkin, Saher S Hammoud, Steven E Domino, Dana C Dolinoy, Jaclyn M Goodrich, Rita Loch-Caruso, Vasantha Padmanabhan, Kelly M Bakulski

## Abstract

The placenta mediates adverse pregnancy outcomes, including preeclampsia, which is characterized by gestational hypertension and proteinuria. Placental cell type heterogeneity in preeclampsia is not well-understood and limits mechanistic interpretation of bulk gene expression measures. We generated single-cell RNA-sequencing samples for integration with existing data to create the largest deconvolution reference of 19 fetal and 8 maternal cell types from placental villous tissue at term (n=15,532 cells). We deconvoluted eight published microarray case-control studies of preeclampsia (n=330). Deconvolution revealed excess extravillous trophoblasts and fewer mesenchymal cells. Adjustment for cellular composition reduced preeclampsia-associated differentially expressed genes (FDR<0.05) from 1,224 to 0, whereas pathway alterations exhibiting a metabolic adaptation to hypoxia were robust to cell type adjustment. Cellular composition explained 35.1% of the association between preeclampsia and *FLT1* overexpression. Our findings indicate substantial placental cellular heterogeneity in preeclampsia that predicts previously observed bulk gene expression differences. Our deconvolution reference lays the groundwork for cellular heterogeneity-aware investigation into placental dysfunction and adverse birth outcomes.

## Introduction

The public health burden of adverse pregnancy outcomes is substantial. An important example is preeclampsia, which affected 6.5% of all pregnant people in the United States in 2017 and is characterized by high maternal blood pressure and damage to other organ systems. Adverse pregnancy outcomes may lead to myriad health complications including elevated risk of chronic diseases throughout the life course [1]. The placenta, a temporary organ that develops early in pregnancy, promotes maternal uterine artery remodeling; mediates transport of oxygen, nutrients, and waste [2]; secretes hormones to regulate pregnancy; metabolizes various macromolecules and xenobiotics; and can serve as a selective barrier to some, but not all, pathogens and xenobiotics [3]. The executive summary of the *Placental Origins of Adverse Pregnancy Outcomes: Potential Molecular Target*s workshop recently concluded that most adverse pregnancy outcomes are rooted in placental dysfunction [4]. Despite this, the molecular underpinnings of placental dysfunction are poorly understood.

Placenta-specific cell types, including cytotrophoblasts, syncytiotrophoblasts, extravillous trophoblasts, and placental resident macrophage Hofbauer cells are all essential for placental development, structure, and function [5]. Dysfunction of these specific cell types likely plays a role in placental pathogenesis. For example, extravillous trophoblasts are responsible for invading into the maternal decidua early in pregnancy to remodel uterine arteries and increase blood flow to the placenta [2]. Inadequate or inappropriate invasion of extravillous trophoblasts has previously been implicated in preeclampsia etiology [6–8]. Despite some knowledge of the roles of specific placental cell types in the development of preeclampsia, relatively little is known about how individual cell types contribute to placental dysfunction.

Existing research models used to investigate the function and dysfunction of individual cell types are limited. Protocols to isolate primary placental cells for experimental research are restricted to one or few cell types [9–14]. Cell type-specific assays are costly and require special techniques or training resulting in small sample sizes and have not yet been scalable to large epidemiological studies [15–17]. Furthermore, placental cell lines such as BeWo, derived from choriocarcinoma [18], and HTR-8/SVneo, immortalized by SV40 [19], are typically derived by processes that alter the DNA of the cells, limiting their in vivo translatability. Consequently, the characteristics of even healthy placental cell type function and especially their connections to adverse outcomes such as preeclampsia are incompletely understood.

Measures of gene expression in bulk placental tissue are used to better understand the biological mechanisms underlying adverse pregnancy outcomes [20–22] and are common in epidemiological studies [23]. Gene expression profiles differ systematically by cell type [24, 25]. Thus, bulk placental tissue-level gene expression measurements represent a convolution of gene expression signals from individual cells and cell types [26, 27]. Deconvolution refers to the bioinformatic process of estimating the distribution of cell types that constitute the tissue [28, 29]. Deconvoluting tissue-level gene expression profiles is essential to eliminate effects introduced by unmodeled cell type proportions [30] by disentangling shifts in cell type proportions from direct changes to cellular gene expression [31]. Reference-based deconvolution boasts biologically interpretable cell type proportion estimates with few modeling assumptions but relies on independently collected cell type-specific gene expression profiles as inputs [31]. Prior placental cell type-specific gene expression measures from term villous tissue [16] had a limited number of biological replicates and included neither technical replicates nor benchmarking against physically isolated placental cell types. A robust, accessible, and publicly available gene expression deconvolution reference is currently unavailable for healthy placental villous tissue.

To advance the field of perinatal molecular epidemiology, our goal was to develop an accessible and robust gene expression deconvolution reference for healthy placental villous tissue at term. We generated single-cell RNA-sequencing data with technical replicates for integration with existing cell type-specific placental gene expression data [16]. Additionally, we benchmarked these single-cell cell type-specific gene expression profiles against placental cell types isolated with more conventional fluorescence-activated cell sorting followed by RNA-sequencing. Finally, to apply our deconvolution approach and assess links between preeclampsia and placental cell types and their proportions, we applied our placenta cell type gene expression reference to deconvolute bulk placental tissues in a secondary data analysis of a case-control study [32] of preeclampsia.

## Results

### Single-cell gene expression map of healthy placental villous tissue

From healthy term placental villous tissue, 9,942 cells across a total of two biological replicates and two technical replicates were sequenced and analyzed. These data were combined with single-cell RNA-sequencing data of 6,313 cells from three healthy term villous tissue samples in a previously published study [16] (**Supplementary Table 1**). Cells were excluded if they were doublets or outliers in total RNA content, number of genes detected, or mitochondrial gene expression (**Supplementary Figures 1-2**). Fetal or maternal origin was determined by genetic variation in sequencing data. Fetal sex was determined by *XIST* expression (**Supplementary Figure 3**). The final analytic sample included 15,532 cells and 36,601 genes across five biological replicates, two of which had a technical replicate.

Uniform manifold approximation and projection [33] was used to visualize sequencing results in two dimensions (**Figure 1A**). Cells clustered into 19 fetal and 8 maternal cell types with 66.5% of all cells of fetal origin (**Table 1**). Observed placenta-specific trophoblast cell types included cytotrophoblasts, proliferative cytotrophoblasts, extravillous trophoblasts, and syncytiotrophoblasts. Proliferative cytotrophoblasts were distinguished by overexpression of genes related to the mitotic cell cycle (p_adj_=7.1×10^−64^) (**Supplementary Figure 4**). Other fetal-specific cell types included mesenchymal stem cells, fibroblasts, endothelial cells, and Hofbauer cells (**Figure 1B**). Fibroblasts were distinguished from mesenchymal stem cells by their overexpression of type 1 collagen genes, e.g., *COL1A1* (log_2_-Fold-Change (log_2_FC) = 3.0, p_adj_=2.48×10^−16^). Lymphocytes, B cells, and monocytes were generally of mixed maternal and fetal origin (**Figure 1B-C**).

**Table 1.** Number of cells captured by single-cell RNA sequencing in the final analytic dataset for each cell type by sample source. Overall cell composition by cell count provided for each cell type.

| Cell Type | 1A | 1B | 2A | 2B | 3 | 4 | 5 | Fetal (%) | Maternal (%) |
| --- | --- | --- | --- | --- | --- | --- | --- | --- | --- |
| Fetal B Cells | 111 | 152 | 199 | 233 | 39 | 28 | 16 | 7.5% | - |
| Fetal CD14+ Monocytes | 240 | 196 | 186 | 204 | 52 | 28 | 43 | 9.2% | - |
| Fetal CD8+ Cytotoxic T Cells | 220 | 210 | 148 | 163 | 79 | 76 | 6 | 8.7% | - |
| Fetal Cytotrophoblasts | 4 | 9 | 1 | 1 | 378 | 921 | 891 | 21.4% | - |
| Fetal Endothelial Cells | 5 | 11 | 23 | 15 | 1 | 5 | 6 | 0.6% | - |
| Fetal Extravillous Trophoblasts | 0 | 0 | 0 | 0 | 11 | 229 | 6 | 2.4% | - |
| Fetal Fibroblasts | 1 | 1 | 0 | 3 | 26 | 17 | 50 | 0.9% | - |
| Fetal GZMB+ Natural Killer | 10 | 7 | 7 | 7 | 48 | 16 | 7 | 1.0% | - |
| Fetal GZMK+ Natural Killer | 7 | 13 | 8 | 11 | 107 | 90 | 10 | 2.4% | - |
| Fetal Hofbauer Cells | 26 | 26 | 112 | 134 | 117 | 121 | 277 | 7.9% | - |
| Fetal Memory CD4+ T Cells | 26 | 42 | 43 | 47 | 70 | 54 | 29 | 3.0% | - |
| Fetal Mesenchymal Stem Cells | 54 | 47 | 56 | 65 | 387 | 13 | 512 | 11.0% | - |
| Fetal Naive CD4+ T Cells | 225 | 216 | 227 | 217 | 30 | 56 | 25 | 9.6% | - |
| Fetal Naive CD8+ T Cells | 52 | 38 | 108 | 127 | 17 | 38 | 9 | 3.8% | - |
| Fetal Natural Killer T Cells | 32 | 37 | 24 | 35 | 56 | 43 | 8 | 2.3% | - |
| Fetal Nucleated Red Blood Cells | 1 | 0 | 1 | 0 | 25 | 6 | 15 | 0.5% | - |
| Fetal Plasmacytoid Dendritic Cells | 5 | 4 | 2 | 7 | 24 | 18 | 8 | 0.7% | - |
| Fetal Proliferative Cytotrophoblasts | 0 | 1 | 1 | 0 | 124 | 200 | 304 | 6.1% | - |
| Fetal Syncytiotrophoblasts | 4 | 15 | 3 | 2 | 6 | 80 | 0 | 1.1% | - |
| Maternal B Cells | 117 | 143 | 348 | 339 | 1 | 21 | 0 | - | 18.6% |
| Maternal CD14+ Monocytes | 284 | 298 | 187 | 212 | 22 | 6 | 8 | - | 19.5% |
| Maternal CD8+ Cytotoxic T Cells | 433 | 419 | 225 | 248 | 22 | 38 | 3 | - | 26.7% |
| Maternal FCGR3A+ Monocytes | 120 | 137 | 50 | 69 | 36 | 11 | 84 | - | 9.7% |
| Maternal Naive CD4+ T Cells | 51 | 36 | 47 | 58 | 9 | 19 | 3 | - | 4.3% |
| Maternal Naive CD8+ T Cells | 69 | 94 | 178 | 168 | 6 | 32 | 3 | - | 10.6% |
| Maternal Natural Killer Cells | 110 | 138 | 64 | 59 | 12 | 8 | 3 | - | 7.6% |
| Maternal Plasma Cells | 36 | 37 | 30 | 46 | 0 | 7 | 2 | - | 3.0% |

**Figure 1.**
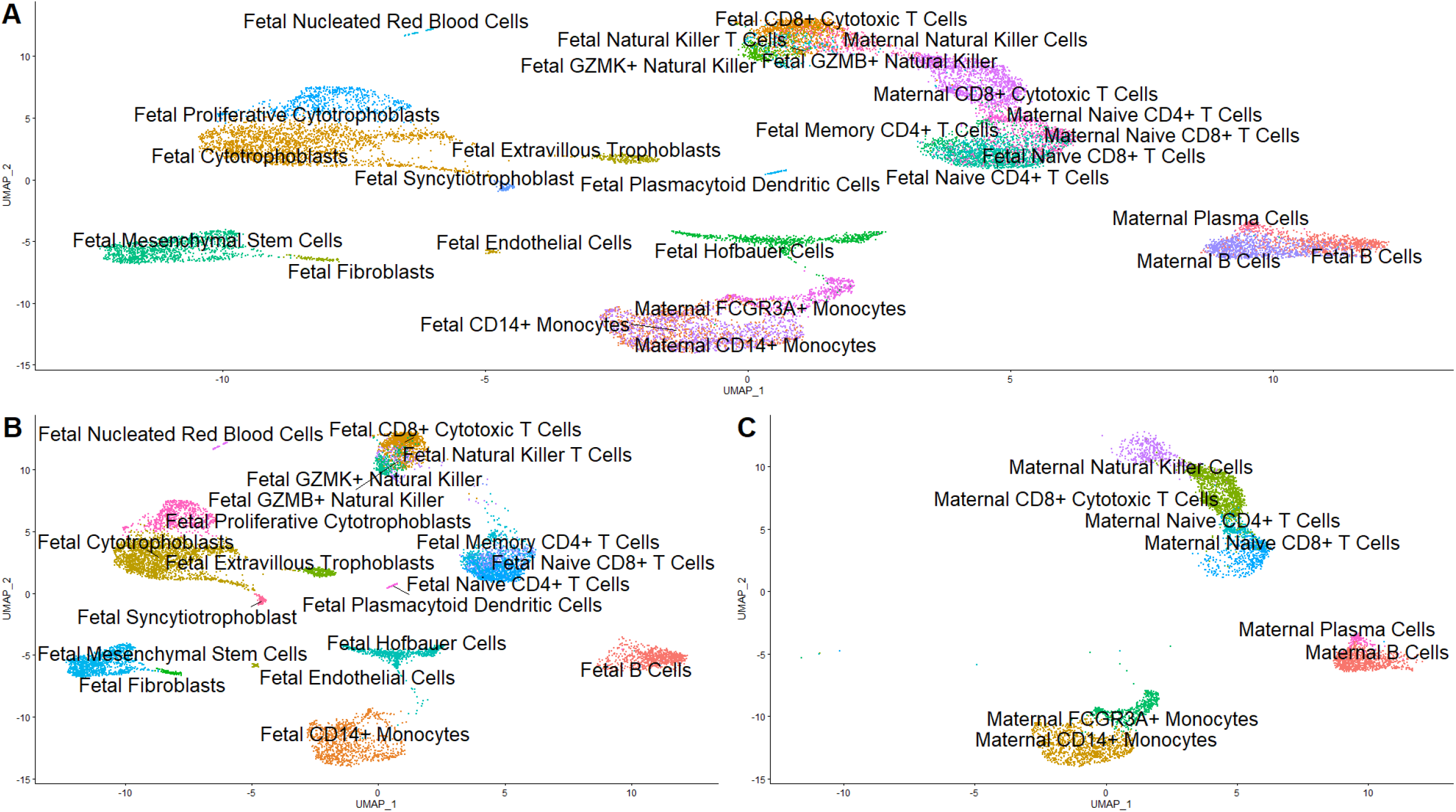
(A) Uniform Manifold Approximation and Projection (UMAP) plot of all cells, with each cell colored by cell type cluster. (B) UMAP plot of fetal cells only, with each cell colored by cell type cluster. (C) UMAP plot of maternal cells only, with each cell colored by cell type cluster.

Cell type-defining transcripts were identified by comparing the expression of a transcript in one cell type against that gene’s average expression across all other cell types (**Supplementary Table 2**). *FLT1* expression was upregulated in extravillous trophoblasts (log_2_FC=3.28), cytotrophoblasts (log_2_FC=0.73), and proliferative cytotrophoblasts (log_2_FC=0.27). Trophoblast cell types had the largest and most diverse transcriptomes, characterized by the largest number of RNA molecules and detected genes per cell (**Supplementary Figure 5**). Cell-type defining biological processes were highlighted with functional enrichment analysis (**Supplementary Table 3**). For example, syncytiotrophoblasts were enriched for transcripts involved in hormone biosynthesis (p_adj_<0.001) and growth hormone receptor signaling (p_adj_=0.003). Technical replication in samples 1 and 2 appeared high in Uniform Manifold Approximation and Projection (UMAP) space (**Supplementary Figure 6A-B**). Indeed, the average intra-cluster gene expression between technical replicates had an average Pearson correlation (mean ± standard deviation) of 0.94 ± 0.14 for sample 1 and 0.88 ± 0.20 for sample 2 (p-values<0.001).

### Fluorescence-activated cell sorting of major placental cell types

We isolated bulk placental villous tissue, enriched syncytiotrophoblasts, and sorted five cell types (Hofbauer cells, endothelial cells, fibroblasts, leukocytes, extravillous trophoblasts, and cytotrophoblasts) via fluorescence-activated cell sorting (FACS) from four healthy term, uncomplicated Cesarean sections for bulk RNA sequencing, labelled samples 1 (same sample source as single-cell RNA sequencing sample 1), 6, 7, and 8 (**Supplementary Table 4**). Principal components analysis of sorted-cell bulk RNA sequencing revealed three loosely defined clusters along principal component (PC) 1: (i) composite tissue and syncytiotrophoblast, (ii) Hofbauer and fibroblast, and (iii) cytotrophoblast, extravillous trophoblast, and endothelial. Leukocyte samples were scattered along PC1 (**Supplementary Figure 8**). Comparison of the expression of a transcript in one cell type against that gene’s average expression in all other cell types identified cell type-defining transcripts (**Figure 2**). 37,929 genes were tested. 746 genes were dropped from the syncytiotrophoblast contrast due to excessively low counts, low variability, or extreme outlier status. Large-scale gene expression differences were observed for each cell type (**Supplementary Table 5**). Functional analysis of cell type-defining transcripts revealed cell type-defining biological processes (**Supplementary Table 6**). For example, syncytiotrophoblasts were enriched for transcripts relevant to endothelium development, mesenchyme development, and vasculogenesis (p_adj_<0.001). Normalized counts between sorted and single-cell RNA-sequencing cell types had an average Spearman correlation of 0.73 ± 0.09 (p<0.001) (**Supplementary Figure 9**).

**Figure 2.**
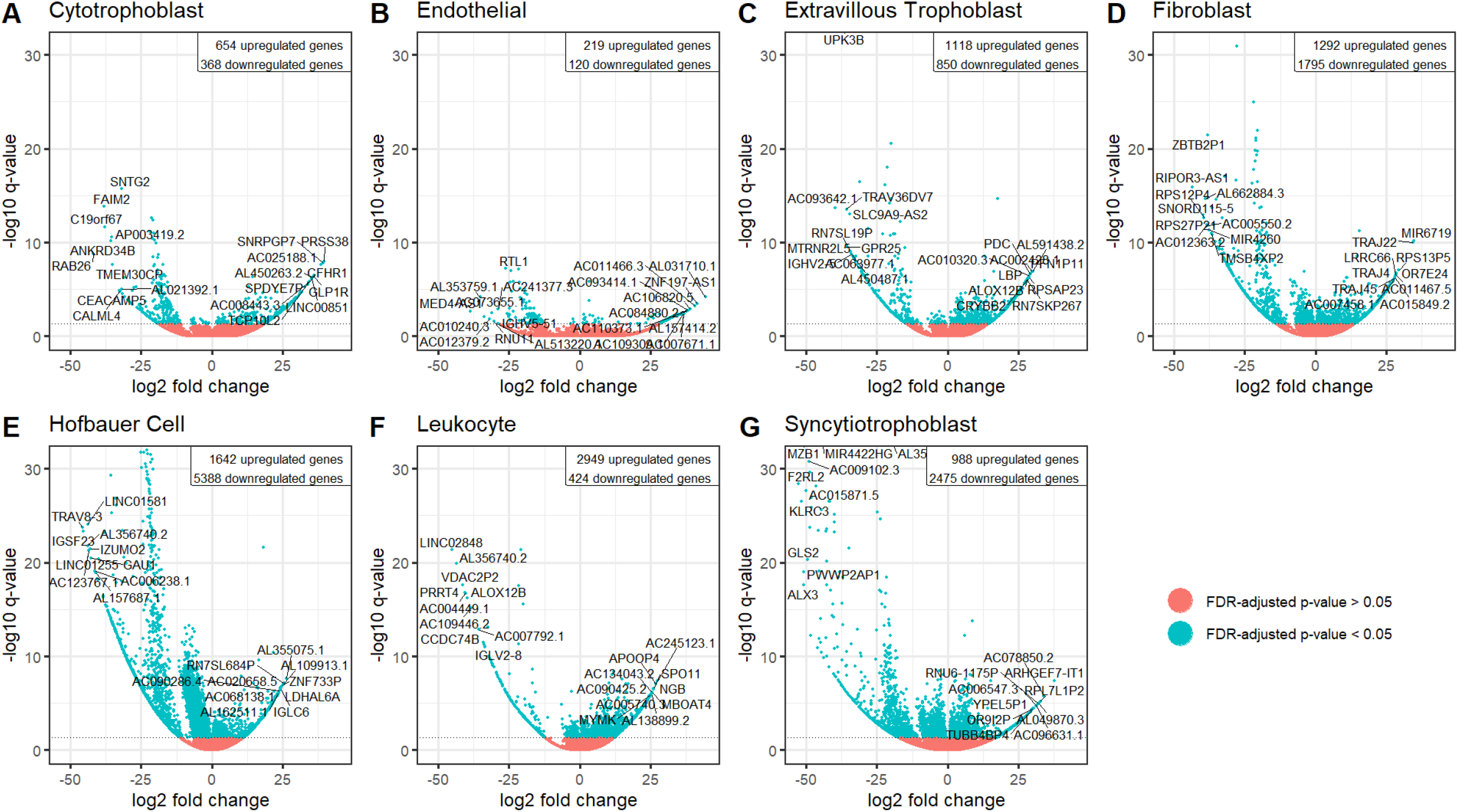
Volcano plots for fluorescence-activated-cell-sorted bulk RNA-seq differential expression in one cell type against average gene expression across other cell types. The y-axis encodes -log10 transformation of the false discovery-controlled q-value, with the cut-off for statistical significance at 0.05. The x-axis encodes log2 fold change of gene expression for the contrast of interest. The upper-right inset describes the number of differentially regulated genes per contrast. 37,929 genes were tested. 746 genes were dropped from the syncytiotrophoblast contrast by DESeq2’s default automatic filtering algorithm due to excessively low counts, low variability, or extreme outlier status. (A) Cytotrophoblast. (B) Endothelial cell. (C) Extravillous trophoblast. (D) Fibroblast. (E) Hofbauer cell. (F) Leukocyte. (G) Syncytiotrophoblast.

### Cell proportion deconvolution of bulk placental tissue dataset

Based on the single cell data, we created a placental signature gene matrix that incorporated an average of 10,044 differentially expressed genes for each of the 27 fetal and maternal cell types (**Supplementary Figure 10**). We applied this signature matrix to estimate cell proportions from bulk placental tissue in 157 preeclampsia cases and 173 controls in a dataset [32] compiled from eight previously published studies [32, 34–40]. Gestational age was 2.2 weeks younger in cases than controls (p-value<0.001, **Table 2**). All deconvoluted samples exhibited high goodness-of-fit between original bulk mixtures and the estimated cell type proportion mixtures (p-values<0.001). Among the signature genes, original bulk and estimated mixtures had a Pearson correlation of 0.71 ± 0.03 and root mean square error of 0.74 ± 0.03 (**Supplementary Table 7**). Fetal naïve CD4+ T cells and fetal B cells were estimated to be at 0% abundance in all samples and were dropped from downstream analyses. Cytotrophoblasts were the most abundant estimated fetal cell type (29% ± 4%) followed by syncytiotrophoblasts (21% ± 4%) and naïve CD8+ T cells (9% ± 2%). The most common maternal cell types were naïve CD8+ T cells (3% ± 2%), natural killer cells (3% ± 2%), and plasma cells (3% ± 2%).

**Table 2.** Demographic characteristics of eight previously published bulk microarray placental gene expression case-control studies (accessed through GSE75010) for deconvolution application testing.

|  | Control<br>(N=173) | Preeclampsia<br>(N=157) | P-value |
| --- | --- | --- | --- |
| Fetal Sex |  |  |  |
| Female | 78 (45.1%) | 81 (51.6%) | 0.28 |
| Male | 95 (54.9%) | 76 (48.4%) |  |
| Gestational Age (wks) |  |  |  |
| Mean (SD) | 35.2 (3.97) | 33.0 (3.17) | <0.001 |
| Median [Min, Max] | 37.0 [25.0, 41.0] | 33.0 [25.0, 39.0] |  |
| Study |  |  |  |
| GSE10588 | 26 (15.0%) | 17 (10.8%) | 0.39 |
| GSE24129 | 8 (4.6%) | 8 (5.1%) |  |
| GSE25906 | 37 (21.4%) | 23 (14.6%) |  |
| GSE30186 | 6 (3.5%) | 6 (3.8%) |  |
| GSE43942 | 7 (4.0%) | 5 (3.2%) |  |
| GSE44711 | 8 (4.6%) | 8 (5.1%) |  |
| GSE4707 | 4 (2.3%) | 10 (6.4%) |  |
| GSE75010 | 77 (44.5%) | 80 (51.0%) |  |

Bivariate batch with Kruskal-Wallis ANOVA (regular ANOVA homogeneity of variances violated) for continuous variables and Chi-square test for categorical outcomes.

### Differentially abundant cell type proportions in preeclampsia cases versus controls

To test for differences in cell proportions between preeclampsia cases and controls (**Supplementary Figure 11**), we fit beta regression models adjusted for each cell type proportion and study source, fetal sex, and gestational age. Fetal memory CD4+ T cells (p=0.03) and extravillous trophoblasts (p<0.001) were more abundant (**Figure 3**) in preeclampsia cases relative to controls. The unadjusted median extravillous trophoblast abundance was 5.3% among cases compared to 2.6% among controls. Mesenchymal stem cells (median percent composition in cases vs. controls, 4.9% vs. 6.2%), Hofbauer cells (6.4% vs. 7.9%), and fibroblasts (6.1% vs. 6.7%) were all less abundant among preeclampsia cases compared to controls (p<0.001). Among maternal cell types, natural killer cells (1.6% vs. 1.4%) were more abundant among preeclampsia cases compared to controls (p=0.002).

**Figure 3.**
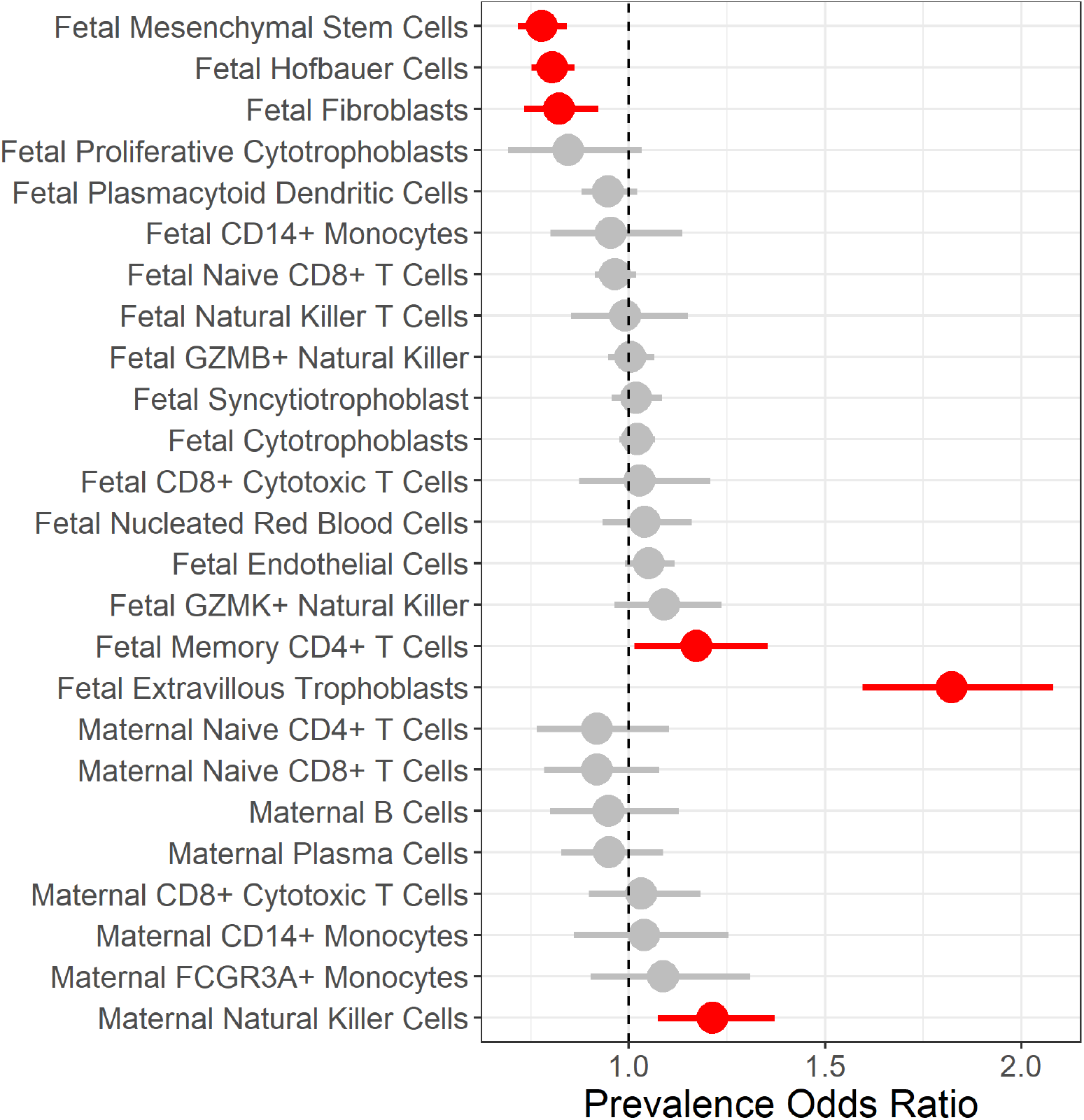
Forest plot of multivariate beta regression models’ prevalence odds ratio adjusted for study source, gestational age, and fetal sex tested for a difference in each cell type’s proportions in cases versus controls. Horizontal lines indicate the range of the 95% confidence interval.

### Differential expression between preeclampsia cases and controls attenuated by cell type proportion adjustment

To test whether microarray gene expression differences between preeclampsia cases and controls are partly driven by differences in cell type abundances, we fit differential gene expression models adjusted for covariates study source, fetal sex, and gestational age with and without adjustment for imputed cell type proportions. To reduce the number of model covariates and account for dependence between deconvoluted cell type proportions, we applied principal components analysis to the imputed cell type proportions. The first five PCs accounted for 77.8% of the variance and were added as additional covariates to form the cell type-adjusted model. Variation in PCs 1 and 2 was largely driven by syncytiotrophoblasts (53.3%) and cytotrophoblast (21.5%) proportions and did not readily separate cases from controls (**Supplementary Figures 12A, C**). Variation in PC3 was largely driven by extravillous trophoblast proportions (69.1%) and to a lesser extent Hofbauer cells (13.5%) and mesenchymal stems cells (6.1%). PC3 provided some separation between cases and controls (**Supplementary Figures 12B, D**).

In the cell type-naïve base model, 1,224 genes were differentially expressed in preeclampsia cases versus controls (**Figure 4A**). Gene set enrichment analysis identified 81 overrepresented pathways in the base model (**Figure 5A**). Biological process pathways such as cellular respiration (q-value<0.001), translational termination (q<0.001), and related pathways were downregulated whereas cell type differentiation pathways such as cornification (q<0.001) and endothelial cell development (q=0.04) were upregulated. Remarkably, when the base model was additionally adjusted for the first five PCs of imputed cell type proportions, there were zero differentially expressed genes between preeclampsia cases and controls (**Figure 4B**). Of the cell type-adjusted results, 40 pathways were overrepresented (**Figure 5B**). Downregulation of translation termination (q<0.001), cellular respiration (q<0.001), and related pathways and upregulation of glycolytic process through fructose 6-phosphate (q=0.01) were robust to cell type proportion adjustment.

**Figure 4.**
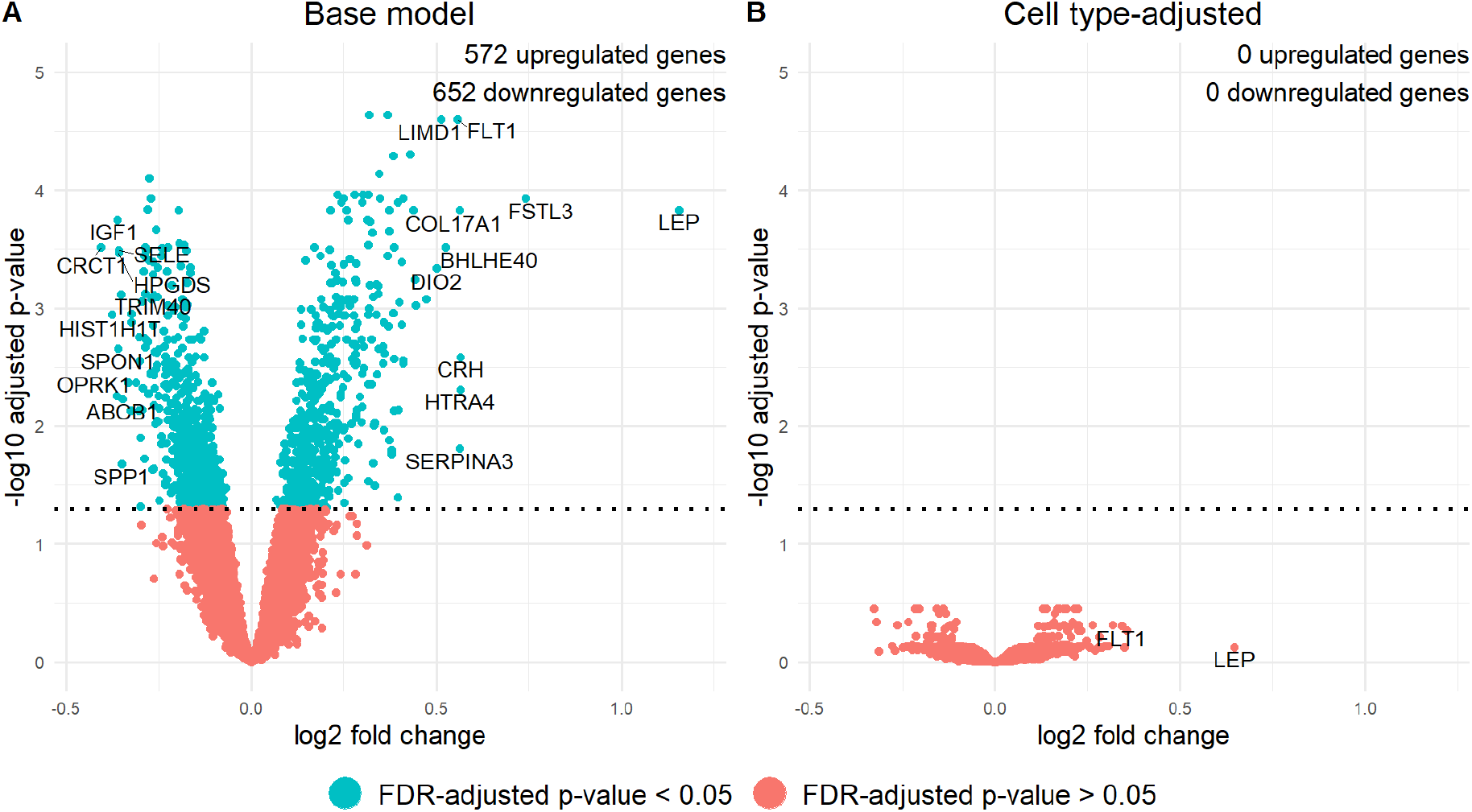
Volcano plots comparing differentially expressed genes in samples from preeclampsia cases versus healthy controls identified as significant by two models: (**A**) the base model adjusted for covariates fetal sex, study source, and gestational age and (**B**) the model additionally adjusted for the first five principal components of estimated cell type proportions. Dotted line represents a false discovery rate-adjusted q-value of 0.05. *FLT1* and *LEP* are labelled as a gene of interest in preeclampsia or as an outlier in log2 fold change, respectively.

**Figure 5.**
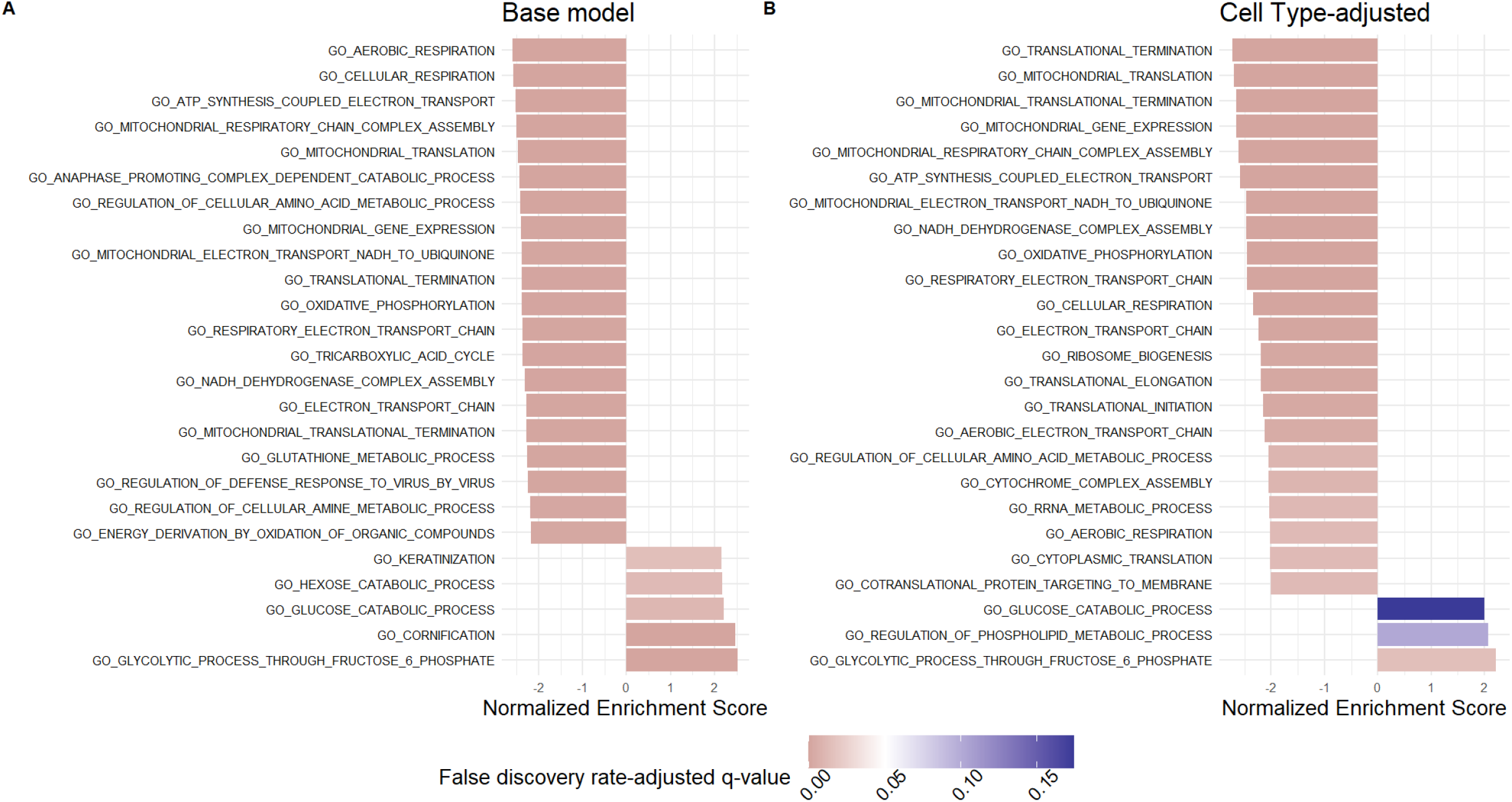
Top Gene Set Enrichment Analysis pathways from the Gene Ontology: Biological Processes database results for the differential expression analysis by preeclampsia case-control status. Results arranged by descending magnitude of the absolute value of the normalized enrichment score. Pathways colored red are significant at a false discovery rate-adjusted (FDR) q-value of 0.05 whereas pathways in blue are statistically insignificant. (**A**) Top pathways from the cell type-unadjusted analysis. (**B**) Top pathways from the cell type-adjusted analysis

### Differential expression of preeclampsia-associated gene *FLT1* mediated by placental cell type proportions

Overexpression of *FLT1* in placental tissue [41–44], detection of a soluble isoform of *FLT1* in maternal circulation [45, 46], and fetal genetic variants near *FLT1* [47] have implicated *FLT1* in preeclampsia etiology. Because we observed cell type-specific expression patterns of *FLT1* in trophoblasts, particularly in extravillous trophoblasts, we hypothesized that the observed attenuation of *FLT1* differential expression may be due in part to the differences in cell type proportions observed between preeclampsia cases and controls. To test this hypothesis, we applied a unified mediation and interaction analysis to quantify the proportion of *FLT1* expression differences mediated by deconvoluted cell type proportions. We did not observe an interaction between preeclampsia status and placental cell composition (average mediated effect percent difference = - 3.8%, 95% CI [-23.4%, 15.7%]). 35.1% (95% CI [25.5%, 46.3%]) of the association between preeclampsia and *FLT1* expression was attributable to differences in placental cell composition between preeclampsia cases and controls (**Figure 6**).

**Figure 6.**
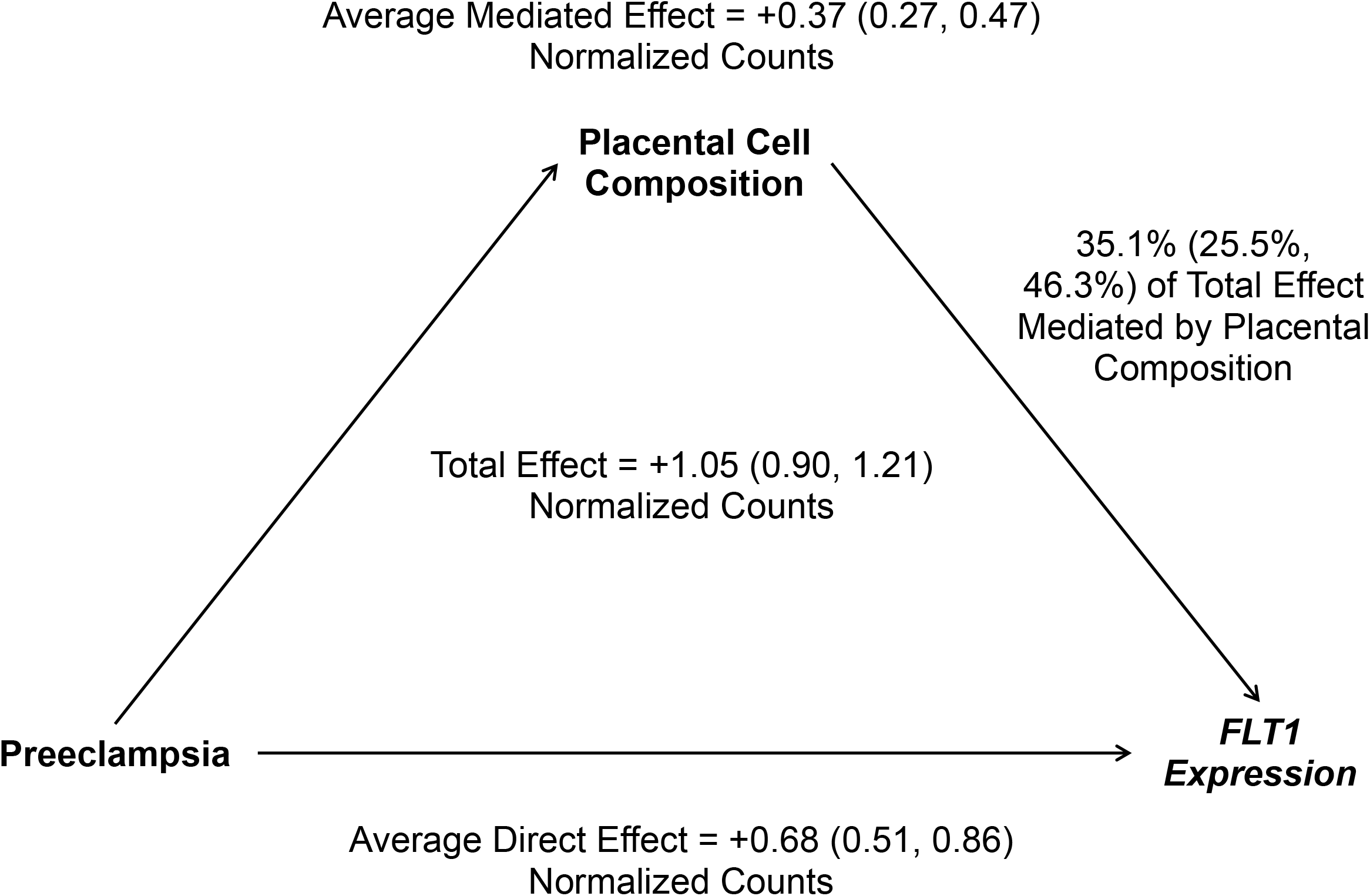
Mediation by placental cell type composition. Placental cell composition was operationalized as first five principal components of estimated cell type proportions. 95% confidence intervals are provided after effect estimates for each model parameter.

## Discussion

To create the largest, publicly available deconvolution reference of 19 fetal and 8 maternal cell type-specific gene expression profiles, we newly sequenced placental villous cells, integrated those results with data from a previously published study, and built a signature gene matrix for deconvolution. To assess reproducibility, we assayed single-cell gene expression profiles for two term placentas in technical replicates following cryopreservation of dissociated villous tissue. To validate single-cell placental cell type expression profiles, we created a novel fluorescence-activated cell sorting scheme to enrich and sequence RNA from five important placental cell types as well as syncytiotrophoblasts. We applied our results to deconvolute cell type proportions in a previously published epidemiologic microarray study of the pregnancy complication preeclampsia, revealing placental cell type proportion differences between preeclampsia cases and controls at term. We then showed that large gene expression differences between preeclampsia cases and controls (n=1,224 genes, p_adj_<0.05) were markedly attenuated after adjustment for cell type proportions (n=0 genes, p_adj_<0.05). Preeclampsia associated pathways that were robust to cell type proportion adjustment included downregulation of replication-related and cellular respiration pathways and upregulation of glycolysis through fructose 6-phosphate. Finally, to quantify the attenuation of differential expression of the preeclampsia biomarker *FLT1*, we applied mediation analysis to show that approximately 35% of the association between increased *FLT1* expression and preeclampsia was attributable to placental cell composition. Cell type proportions may be an important factor in gene expression differences in placental tissue studies.

By integrating our new single-cell RNA-sequencing results with those from a previously published study, our integrated dataset, to our knowledge, is the largest for healthy, term placental villous tissue to date. We document term cell type-specific gene expression patterns for well-characterized placental cell types, including syncytiotrophoblasts [9], cytotrophoblasts [12], and extravillous trophoblasts [13]. In addition, we provide gene expression markers for relatively understudied placental cell types such as endothelial cells, mesenchymal stem cells, and Hofbauer cells as well as maternal peripheral mononuclear cells recovered from the maternal-fetal interface. Compared to the previous analysis of the published samples [16] which relied on predominately sex-specific gene expression markers to differentiate proliferative from non-proliferative cytotrophoblasts, we show that functional enrichment analysis revealed broad upregulation of proliferation pathways in proliferative cytotrophoblasts. We replicated the findings of a previous study [9] that size exclusion filtration can enrich syncytiotrophoblasts from gently digested villous tissue based on comparison to single-cell results. We developed a novel fluorescence-activated cell sorting scheme to simultaneously isolate viable cells from five major placental cell types. This protocol can be used to simultaneously isolate placental cell types for functional assays or other experiments. The low representation of some cell types such as trophoblasts in our single-cell RNA-sequencing results suggest that these cell types may be sensitive to a conventional cryopreservation protocol, an extension of dissociation bias that has been previously documented in other tissues [48]. Future studies may propose alternative approaches to perform unbiased single-cell RNA in placental tissues.

Our preeclampsia findings are consistent with prior pathophysiological understanding of the disorder, and we provide linked cell type and gene expression data in bulk tissue for the first time. Among preeclampsia cases, we observed an elevated proportion of extravillous trophoblasts and underrepresentation of stromal cell types, which may reflect an arrest in placental cell type differentiation and maturation following insufficient uterine spiral artery remodeling implicated in preeclampsia [49–51]. In the cell type-naïve differential expression model, consistent with previous findings, placentas from pregnancies with preeclampsia overexpressed *FLT1, LEP*, and *ENG* [41–44]. In our cell type-adjusted model, *FLT1* and *LEP* remained only nominally significant whereas *ENG* was not differentially expressed. Mediation analysis confirmed that a significant proportion of *FLT1* overexpression was attributable to changes in the cellular composition of the placenta. These results suggest that placental cell type proportion differences may be an overlooked factor in explaining the well-documented association between preeclampsia and *FLT1* expression [41–44]. Downregulation of replication-related and cellular respiration pathways and upregulation of glycolysis through fructose 6-phosphate was robust to cell type adjustment, suggesting intracellular changes to these pathways. Together, these enrichment results suggest a metabolic adaption to hypoxia. Placental hypoxia is characteristic of preeclampsia [52, 53]. Because oxygen tension is a critical factor in trophoblast differentiation, inappropriate oxygenation may partially explain the elevated proportion of extravillous trophoblasts, though regulators of this process such as *HIF1A* and *TGFB3* were not differentially expressed at the tissue level [54]. A recent single-cell RNA-sequencing case-control study of preeclampsia, however, identified upregulation of *TGFB1* in extravillous trophoblasts, potentially indicative of altered trophoblast differentiation or invasion [55, 56]. Consistent with our other findings, this study also observed a similar trend in cell type proportion differences and upregulation of *FLT1* in extravillous trophoblasts and *ENG* in syncytiotrophoblasts among between preeclampsia cases and controls [55]. Future work should consider and account for the cell type-specific expression patterns of genes that regulate placental development or are associated with preeclampsia to better understand preeclampsia etiology.

This study has several strengths. This is one of the first studies to profile the parenchymal healthy term villous tissue in the placenta and we integrate our dataset with samples from a previously published study to generate the largest cell type-specific placental villous tissue gene expression reference to date. Single-cell RNA-sequencing allows us to agnostically capture diverse placental cell types without *a priori* knowledge of cell types and their characteristics. To our knowledge, this is also the first study to demonstrate technical replication of single-cell RNA-sequencing in placental villous tissue. We verify our results with conventional RNA-sequencing of FACS-sorted placental cell types. We confirmed enrichment of syncytiotrophoblasts through size exclusion filtration is consistent with single-cell results. We also applied our findings to a large target deconvolution dataset of preeclampsia that contained placental measures from hundreds of participants across eight different studies. Most importantly, this is the first study of preeclampsia to account for cell type heterogeneity, a critical factor in bulk tissue assays, in an epidemiologic sample.

This study also has several limitations. Although our cellular sample size comprised of 15,532 cells is relatively large compared to previous single-cell RNA-sequencing studies of term placental villous tissue, this dataset still represents a limited biologic replicate sample size. Our newly sequenced samples came from a convenience sample without available demographic information beyond healthy Cesarean-section status. Similarly, the sample size of FACS-sorted tissues was limited, and some cell type fractions were excluded due to low RNA quality. This study did not include placental tissues for single cell analysis from preeclamptic patients to confirm intracellular gene expression changes. Though deconvolution fit statistics were highly significant, we had no gold standard to verify accurate deconvolution of cell counts. Future studies may verify whether cell type proportions estimated in diseased or vaginally delivered tissues are robust to a deconvolution reference generated from healthy villous tissue delivered via Cesarean-section. Residual confounding may remain in our statistical models due to the limited number of common covariates across all eight preeclampsia case-control studies. Due to the nature of villous tissue sampling, our study design is cross-sectional, limiting our ability to establish temporality between exposure and outcome to rule out reverse causation. As with any study conditioned on live birth, selection bias may affect our results. However, the effects of harmful exposures that lead to selection tend to be underestimated in these scenarios [57, 58]. Therefore, our results likely represent a conservative underestimate of the effects of preeclampsia on inappropriate cell composition and preeclampsia status on *FLT1* expression.

In summary, we provide a cell type-specific deconvolution reference via single-cell RNA-sequencing in the parenchymal placental term villous tissue. We verified this reference by developing a novel FACS scheme to characterize six major placental cell types with RNA-sequencing. We applied this deconvolution reference to an epidemiologic preeclampsia dataset to reveal biologically relevant shifts in placental cell type proportions between preeclampsia cases and controls. Once cell type proportion differences were accounted for, differential gene expression differences were markedly attenuated between preeclampsia cases and controls. Enrichment analysis revealed downregulation of replication-related and cellular respiration pathways and upregulation of glycolysis through fructose 6-phosphate in preeclampsia. A substantial proportion of the overexpression of the *FLT1* in preeclampsia was mediated by placental cell composition. This is the first study to evaluate cell type proportion differences in an epidemiological study of placental parenchymal tissue and preeclampsia, or genome-wide gene expression differences adjusting for cell type heterogeneity. These results add to the growing body of literature that emphasizes the centrality of cell type heterogeneity in molecular measures of bulk tissues. We provide a publicly available placental cell type-specific gene expression reference for term placental villous tissue to overcome this critical limitation.

## Methods

### Placental tissue collection and dissociation

Placentas were collected shortly after delivery from healthy, full term, singleton uncomplicated Cesarean sections at the University of Michigan Von Voigtlander Women’s Hospital. Villous placental tissue biopsies were collected and minced for dissociation after cutting away the basal and chorionic plates and scraping villous tissue from blood vessels [12]. We subjected approximately 1g minced dissected villous tissue to the Miltenyi Tumor Dissociation Kit on the GentleMACS Octo Dissociator with Heaters (Miltenyi Biotec) to yield single-cell suspensions of viable placental cells in 5μM StemMACS™ Y27632 (Miltenyi Biotec) in RPMI 1640 (Gibco) according to manufacturer’s instructions for “soft” tumor type. Red blood cells were depleted using RBC lysis buffer (Biolegend) according to manufacturer’s protocol A. Single-cell suspensions were size-filtered at 100μm and subsequently 40μm. To collect a syncytiotrophoblast-enriched fraction, the fraction between 40μm and 100μm was washed from the 40μm strainers [9]. Single-cell suspensions <40μm were cryogenically stored in 5μM StemMACS™ Y27632 90% heat-inactivated fetal bovine serum (Gibco)/10% dimethyl sulfoxide (Invitrogen). For each placenta, additional whole villous tissue samples were stored in RNALater (Qiagen).

### Placental single-cell RNA sequencing

Villous tissue single-cell suspensions were thawed and sorted via fluorescence-activated cell sorting with LIVE/DEAD Near-IR stain (Invitrogen) for viability and forward-scatter and side-scatter profiles to eliminate cellular debris and cell doublets. Viability- and size-sorted single-cell suspensions were submitted to the University of Michigan Advanced Genomics Core for single-cell RNA sequencing. Single cells were barcoded, and cDNA libraries constructed on the Chromium platform (10X Genomics, Single Cell 3’ v2 chemistry). Asymmetric paired-end 110 base pair reads were sequenced on NovaSeq 6000 (Illumina).

### Single-cell RNA-sequencing preprocessing

Raw reads were processed, deconvoluted, droplet filtered, and aligned at the gene level with the Cell Ranger pipeline using default settings (v4.0.0, 10X Genomics) based on the GRCh38 GENCODEv32/Ensembl 98 reference transcriptome with STAR v2.5.1b [59]. Previously published single-cell RNA-sequencing raw data of healthy, term placental villous tissue samples (dbGaP ID phs001886.v1.p1) SRR10166478, SRR10166481, and SRR10166484 downloaded through the NCBI Sequence Read Archive were processed identically [16]. The *freemuxlet* function in the latest version (accessed 2021/12/05) of the ‘popscle’ package was used to assign fetal or maternal origin and identify 120 mosaic doublets for removal based on single nucleotide polymorphisms with minor allele frequency greater than 10% from the 1000 Genomes Phase 3 reference panel (released 2013/05/02) [60]. Per cell quality control criteria were total RNA molecules, unique genes, and percentage of reads mapping to mitochondrial genes [61]. We excluded 1,398 low-quality outlier cells defined as cells that exceeded four median absolute deviations in samples 1 and 2 or three median absolute deviations in samples 3, 4, and 5 using the *quickQCPerCell* function in ‘scater’ (R package, version 1.18.6) with default settings [62] (**Supplementary Figures 1-2**).

### Single-cell RNA-sequencing clustering and cluster annotation

Maternal and fetal cells were split into separate datasets for clustering. Mutual nearest neighbor batch correction by biological replicate with default settings was used to identify cell type clusters and visualize clustering results via uniform manifold projection [33] using *FastMNN* from ‘SeuratWrapper’ (R package, version 0.3.0) [63]. Iterative clustering and sub-clustering with ‘Seurat’ (R package, version 4.0.1) function *FindClusters* at different resolution parameters were evaluated using cluster stability via clustering trees in ‘clustree’ [64, 65]. *A priori* canonical cell type marker gene expression patterns and cluster marker genes were used to assign cell types to cell clusters [16, 66–71]. Cells that fell outside cell type clusters and outlying in doublet density calculated with *computeDoubletDensity* were removed as putative doublets and doublet clusters were identified with *findDoubletClusters* for removal in ‘scDblFinder’ (R package, version 1.4.0) [72]. 723 maternal-maternal or fetal-fetal putative doublets were excluded after integration and clustering. Fetal sex in phs001886.v1.p1 samples was determined by annotation and confirmed with *XIST* expression. The final analytic sample had 15,532 cells and 36,601 genes across five biological and two technical replicates.

### Single-cell RNA-sequencing differential expression and biological pathway enrichment statistical analysis

Technical correlation was assessed by Pearson correlation after averaging the normalized expression for each gene by cluster and by technical replicate. Cluster marker genes were identified in ‘Seurat’ with the *FindAllMarkers* function with default settings on uncorrected single-cell gene expression counts [61, 65]. Specifically, including both maternal and fetal cell types, the expression level in each cell type cluster was compared against the average expression of that gene across all other cell types using the two-tailed Wilcoxon Rank Sum test with significance defined at a false discovery rate-adjusted p-value less than 0.05. Pairwise cluster markers were identified in ‘Seurat’ with the *FindMarkers* function with an identical testing regime. Overexpressed genes were ranked by decreasing log-fold change for functional enrichment analysis with ‘gprofiler2’ (R package, version 0.2.0, database version e102_eg49_p15_7a9b4d6) using annotated genes as the universe, excluding electronically generated annotations, with the g:SCS multiple testing correction method applying significance threshold 0.05 [73].

### Fluorescence-activated cell sorting of major placental cell types from villous tissue

Villous tissue single-cell suspensions were quickly thawed and stained with 5 fluorescently labeled antibodies (CD9-FITC, CD45-APC, HLA-A,B,C-PE/Cy7, CD31-BV421, and HLA-G-PE) as well as the LIVE/DEAD Near-IR stain (Invitrogen) to isolate 6 viable populations of placental cells by fluorescence activated cell sorting at the University of Michigan Flow Cytometry Core Facility. Initial flow cytometry experiments included fluorescence minus one, single color compensation, and isotype controls. Isotype controls were found to be the most conservative and were consequently included in all sorting experiments, as well as single-color compensation controls due to the large number of colors used in sorting. The six populations of cells were Hofbauer cells, endothelial cells, fibroblasts, leukocytes, extravillous trophoblasts, and cytotrophoblasts We developed a novel five-marker cell surface fluorescence activated cell sorting (FACS) scheme to sort cytotrophoblasts (HLA A,B,C-), endothelial cells (CD31+), extravillous trophoblasts (HLA-G+), fibroblasts (CD9+), Hofbauer cells (CD9-), and leukocytes (CD45+/CD9+) from villous tissue (**Supplementary Figure 6**) [10, 11, 13, 74–81]. Syncytiotrophoblast fragments were enriched from villous tissue digests. We isolated cell type fractions and composite villous tissue from four healthy term, uncomplicated Cesarean sections, labelled samples 1 (same sample source as single-cell RNA-sequencing sample 1), 6, 7, and 8. We subjected 24 cell type fractions with sufficient RNA content to RNA-sequencing, including five composite, two cytotrophoblast, one endothelial, three extravillous trophoblast, three fibroblast, four Hofbauer cell, four leukocyte, and two syncytiotrophoblast fractions (**Supplementary Table 4**).

Detailed antibody information: FITC, marker CD9: Mouse IgG1-kappa, clone HI9a, Biolegend #312103, lot B188319, Biolegend #312104, lot B232916; isotype control: clone MOPC-21 Biolegend #400107, Lot B199152. APC, marker CD45: Mouse IgG1-kappa, clone 2D1, Biolegend #368511, Lot B215062; isotype control: clone MOPC-21, Biolegend #400121, lot B216780. PE/CY-7, marker HLA-ABC: Mouse IgG2a-kappa, clone W6/32, Biolegend #311429, lot B188649, Biolegend #3111430, lot B238602; isotype control: clone MOPC-173, Biolegend #400231, lot B209000. BV421, marker CD31: Mouse IgG1-kappa, clone WM59, Biolegend #303123, lot B204347, Biolegend #303124, lot B232010; isotype control: clone MOPC-21, Biolegend #400157, lot B225357. PE, marker HLA-G: Mouse IgG2a-kappa, clone 87G, Biolegend #335905, lot B222326, Biolegend #335906, lot B199294; isotype control clone MOPC-173, Biolegend #400211, lot B227641. Mouse IgG1-kappa, clone MEM-G/9, Abcam #24384 Lot GR3176304-1; isotype control: monoclonal, Abcam #ab81200, lot GR267131-1. Validation information available on manufacturer’s website under the catalog ID for each antibody.

A cut-off of 0.1% events was used to set a series of gates. Cells were first gated on size and granularity (FSC-HxSSC-H) to eliminate debris (not shown), followed by doublet discrimination (FSC-HxFSC-W and SSC-HxSSC-W) (not shown). Ax750 was used to sort on viability (not shown). Extravillous trophoblasts were isolated based on Human Leukocyte Antigen-G (HLA-G) expression (**Supplementary Figure 7A**). Cytotrophoblasts are HLA-ABC negative (**Supplementary Figure 7B**). HLA-ABC positive cells were then subjected to a CD45/CD9 gate to isolate Hofbauer cells and a heterogeneous population of leukocytes (**Supplementary Figure 7C**). Finally, CD45-/CD9-population is sorted into the endothelial or fibroblast bins based on CD31 expression (**Supplementary Figure 7D**).

### Bulk placental tissue and sorted placental cell type RNA extraction and sequencing

Approximately 2mg of bulk RNALater-stabilized (Qiagen) bulk villous tissue was added to 350μL 1% β-mercaptoethanol (Sigma-Aldrich) RLT Buffer Plus (Qiagen) to Lysing Matrix D vials (MP Biomedicals). Samples were disrupted and homogenized on the MP-24 FastPrep homogenizer (MP Biomedicals) at 6m/s, setting MP24×2 for 35s. For the homogenized bulk villous tissue, syncytiotrophoblast-enriched fraction, and sorted cell types, RNA extraction was completed according to manufacturer’s instructions using the AllPrep DNA/RNA Mini Kit (Qiagen) and stored at -80°C. RNA samples were submitted to the University of Michigan Advanced Genomics Core for RNA sequencing. Ribosomal RNAs were depleted with RiboGone (Takara) and libraries were prepared with the SMARTer Stranded RNA-Seq v2 kit (Takara). Paired- or single-end 50 base pair reads were sequenced on the HiSeq platform (Illumina). Raw RNA reads were assessed for sequencing quality using ‘FastQC’ v0.11.5 [82] and ‘MultiQC’ v1.7 [83]. Reads were aligned to the GRCh38.p12/ GENCODEv28 reference transcriptome using ‘STAR’ v2.6.0c with default settings [59]. featureCounts from ‘subread’ v1.6.1 was used to quantify and summarize gene expression with default settings [84].

### Validation testing: Sorted placental cell type differential expression analysis and comparison to single-cell results

Cell type-defining transcripts were identified using the negative binomial linear model two-tailed Wald test in ‘DESeq2’ (R package, version 1.30.1) adjusted for biological replicate using default settings with contrasts comparing the expression of a gene in one cell type against the average expression across all other cell types at a false discovery rate-adjusted p-value less than 0.05 [85]. Differentially expressed genes for each contrast were descending-ranked by absolute value of the test statistic for gene set enrichment analysis in desktop version GSEA 4.1.0 with the GSEAPreranked tool with default settings [86, 87]. Sorted- and single-cell read counts were appropriately library-normalized and log-transformed. Diverse fetal and maternal immune cell types from the single-cell RNA-sequencing data were summed and collapsed to one leukocyte category, cytotrophoblast subtypes to one cytotrophoblast category, and mesenchymal stem cells and fibroblasts to one fibroblast category. Normalized sorted cell type gene expression was compared to normalized single-cell RNA-sequencing results using Pearson and Spearman correlation coefficients to validate single-cell expression results.

### Application testing: Bulk placenta gene expression dataset and CIBERSORTx deconvolution

Bulk placental tissue microarray gene expression (previously batch-corrected and normalized) from eight preeclampsia case-control studies was downloaded from the NCBI Gene Expression Omnibus (accession number GSE75010) for deconvolution [32]. We used the online version of CIBERSORTx (cibersortx.standford.edu, accessed 2021-01-25) to create a signature gene expression matrix for deconvolution with the following parameters: differential expression q-value<0.01, no minimum gene expression cutoff, and a 2000 gene feature selection ceiling [88]. We used the signature matrix to estimate constituent cell type proportions in GSE75010 using CIBERSORTx with cross-platform S-mode batch correction and 100 permutations to evaluate imputation goodness-of-fit.

### Application testing: Preeclampsia case-control differential cell type abundance, differential gene expression statistical analysis, and mediation analysis

To test for differences in estimated cell type proportions between preeclampsia cases and controls, estimated cell type proportions for GSE75010 were regressed on preeclampsia case-control status using beta regression models adjusted for gestational age, sex, and study source [89]. Statistical significance was assessed using the two-tailed Wald test applying a nominal significance threshold of 0.05. Cell types imputed at zero percent abundance across all samples were excluded.

Differential expression analysis was conducted in *limma* [90] with default linear models adjusted for gestational age, fetal sex, and study source. A cell type-adjusted model was built on the base model additionally adjusted for the first five principal components of deconvoluted cell type proportions. Principal components analysis was performed with *prcomp* from ‘stats-package’ (R, version 4.0.5) without scaling and default settings. Statistical significance was assessed at false discovery rate-adjusted q-value<0.05. Differentially expressed genes were descending-ranked by absolute value of the test statistic for gene set enrichment analysis in desktop version GSEA 4.1.0 with the GSEAPreranked tool with default settings [86, 87].

A unified mediation and interaction analysis [91] was conducted in ‘CMAverse’ (R package, version 0.1.0) [92] via the g-formula approach [93] to estimate causal randomized-intervention analogues of natural direct and indirect effects [94] through direct counterfactual imputation. The model was operationalized with preeclampsia status as the binary exposure, normalized *FLT1* expression as the continuous outcome, and the first five principal components of deconvoluted cell type proportions as continuous mediators. Baseline covariates included fetal sex and study source. Continuous gestational age was included as a confounder of the mediator-outcome relationship affected by the exposure. Confidence intervals were bootstrapped with 1000 boots with otherwise default settings. Statistical tests were two-tailed and interpreted at a p-value significance threshold of 0.05.

### Statistical Information

Technical replication measured by average intra-cluster gene expression between technical replicates was tested via the two-tailed Pearson correlation test within Samples 1 and 2 assessed across all 36,601 genes. The number of cells contributing expression data for each cell type are available in **Table 1**. Single-cell cluster marker genes were identified in ‘Seurat’ with the *FindAllMarkers* function with default settings on uncorrected single-cell gene expression counts [61, 65]. Specifically, including cells from both maternal and fetal cell types, the expression level in each cell type cluster was compared against the average expression of that gene across all other cell types using the two-tailed Wilcoxon Rank Sum test with significance defined at a false discovery rate-adjusted p-value less than 0.05 (n=15,532 cells). Pairwise cluster markers were identified in ‘Seurat’ with the *FindMarkers* function with an identical testing regime (n=2,835 cells for proliferative vs. non-proliferative cytotrophoblasts). Overexpressed genes (498.37 ± 400.82 genes across 27 cell type clusters or 746 differentially expressed genes for proliferative vs. non-proliferative cytotrophoblasts) were ranked by decreasing log-fold change for functional enrichment analysis with ‘gprofiler2’ (R package, version 0.2.0, database version e102_eg49_p15_7a9b4d6) using annotated genes as the universe, excluding electronically generated annotations, with the g:SCS multiple testing correction method applying significance threshold 0.05 [73]. Overexpressed genes per cell type cluster are available in **Supplementary Table 2**.

Cell type-defining transcripts were identified using the negative binomial linear model two-tailed Wald test in ‘DESeq2’ (R package, version 1.30.1) adjusted for biological replicate using default settings with contrasts comparing the expression of a gene in one cell type against the average expression across all other cell types at a false discovery rate-adjusted p-value less than 0.05 [85] (n=19 cell type fraction samples with breakdown by cell type available in **Supplementary Table 4**). Overexpressed genes for each contrast (1,261 ± 866.83 genes across 7 cell types) were descending-ranked by absolute value of the test statistic for gene set enrichment analysis in desktop version GSEA 4.1.0 with the GSEAPreranked tool with default settings [86, 87]. Differentially expressed genes per cell type available in **Supplementary Table 5** and number of differentially expressed genes is summarized in **Figure 2**. Sorted- and single-cell read counts were appropriately library-normalized and log-transformed. Diverse fetal and maternal immune cell types from the single-cell RNA-sequencing data were summed and collapsed to one leukocyte category, cytotrophoblast subtypes to one cytotrophoblast category, and mesenchymal stem cells and fibroblasts to one fibroblast category. Normalized sorted cell type gene expression (n=19 cell type fraction samples) was compared to normalized single-cell RNA-sequencing (n=15,484 cells) results across 7 cell types using two-tailed Pearson and Spearman correlation coefficient tests to validate single-cell expression results across 27,490 common genes (**Supplementary Figure 9**). Statistical tests were interpreted at a p-value significance threshold of 0.05.

Bulk placental tissue microarray gene expression (previously batch-corrected and normalized) from eight preeclampsia case-control studies was downloaded from the NCBI Gene Expression Omnibus (GSE75010) for deconvolution (n=330) [32]. We used the online version of CIBERSORTx (cibersortx.standford.edu, accessed 2021-01-25) to create a signature gene expression matrix for deconvolution with the following parameters: differential expression q-value<0.01, no minimum gene expression cutoff, and a 2000 gene feature selection ceiling [88]. We used the signature matrix to estimate constituent cell type proportions in GSE75010 using CIBERSORTx with cross-platform S-mode batch correction and 100 permutations to evaluate imputation goodness-of-fit.

To test for differences in estimated cell type proportions between preeclampsia cases and controls (n=330), estimated cell type proportions for GSE75010 were regressed on preeclampsia case-control status using beta regression models (n=25 cell type proportion outcomes) adjusted for gestational age, sex, and study source [89]. Cell types imputed at zero percent abundance across all samples were excluded (n=2 excluded). Statistical significance was assessed using the two-tailed Wald test applying a nominal significance threshold of 0.05.

Differential expression analysis was conducted in *limma* [90] with default linear models adjusted for gestational age, fetal sex, and study source (n=330). A cell type-adjusted model was built on the base model additionally adjusted for the first five principal components of deconvoluted cell type proportions. Principal components analysis was performed with *prcomp* from ‘stats-package’ (R, version 4.0.5) without scaling and default settings. Statistical significance was assessed at false discovery rate-adjusted q-value<0.05. Differentially expressed genes were descending-ranked by absolute value of the test statistic for gene set enrichment analysis in desktop version GSEA 4.1.0 with the GSEAPreranked tool with default settings [86, 87].

A unified mediation and interaction analysis [91] was conducted in ‘CMAverse’ (R package, version 0.1.0) [92] via the g-formula approach [93] to estimate causal randomized-intervention analogues of natural direct and indirect effects [94] through direct counterfactual imputation. The model (n=330) was operationalized with preeclampsia status as the binary exposure, normalized *FLT1* expression as the outcome, and the first five principal components of deconvoluted cell type proportions as continuous mediators. Baseline covariates included fetal sex and categorical study source. Continuous gestational age was included as a confounder of the mediator-outcome relationship affected by the exposure. Confidence intervals were bootstrapped with 1000 boots with otherwise default settings. Statistical tests were two-tailed and interpreted at a p-value significance threshold of 0.05.

## Declarations

## Acknowledgements

This research was supported by the Michigan Lifestage Environmental Exposures and Disease center (P30 ES017885), Michigan State University’s Environmental Influences on Child Health Outcomes program (UG3 OD023285, UH3 OD023285), the University of Michigan M-cubed pilot grant program, and the Puerto Rico Testsite for Exploring Contamination Threats (P42 ES017198). K.A.C. was supported by the National Institutes of Health (T32 HG00040). J.A.C. was supported by the Ravitz Family Foundation, the Forbes Institute for Cancer Discovery at the University of Michigan Rogel Cancer Center, and the National Institutes of Health (R01 ES028802, U01 ES026697, and P30 CA046592). M.P. was supported by the National Institutes of Health (T32 ES007062). E.R.E was supported by the National Institutes of Health (T32 DK071212). K.M.B. was supported by research grants from the National Institute of Environmental Health Sciences (R01 ES025531; R01 ES025574). We also acknowledge the submitting investigators of previously published placental single-cell RNA sequencing data (dbGaP accession number phs001886.v1.p1), whose work was sponsored in part by the Perinatology Research Branch, Division of Obstetrics and Maternal-Fetal Medicine, Division of Intramural Research, Eunice Kennedy Shriver National Institute of Child Health and Human Development, National Institutes of Health, U.S. Department of Health and Human Services.

## Author contributions

**Kyle Campbell:** Conceptualization, Methodology, Software, Validation, Formal analysis, Investigation, Data Curation, Writing – Original Draft, Visualization **Justin A. Colacino:** Conceptualization, Methodology, Software, Validation, Investigation, Resources, Writing – Review & Editing, Visualization, Supervision, Project administration, Funding acquisition **Muraly Puttabyatappa:** Investigation, Writing – Review & Editing, Project administration **John Dou:** Software, Validation, Data Curation **Elana R. Elkin:** Writing – Review & Editing **Saher Sue Hammoud:** Methodology, Writing – Review & Editing, Funding acquisition **Steven E. Domino:** Methodology, Resources, Writing – Review & Editing, Project administration, Funding acquisition **Dana C. Dolinoy:** Writing – Review & Editing, Funding acquisition **Jaclyn M. Goodrich:** Writing – Review & Editing, Funding acquisition **Rita Loch-Caruso:** Conceptualization, Methodology, Resources, Writing – Review & Editing, Supervision, Project administration, Funding acquisition **Vasantha Padmanabhan:** Conceptualization, Resources, Writing – Review & Editing, Project administration, Funding acquisition **Kelly M. Bakulski:** Conceptualization, Methodology, Software, Validation, Formal analysis, Investigation, Resources, Data Curation, Writing – Original Draft, Visualization, Supervision, Project administration, Funding acquisition

## Human Subject Data

This study did not collect data on human subjects. Study protocols for tissue collection were approved by the University of Michigan Institutional Review Board (HUM00017941, HUM00102038).

## Data Availability

The cell type signature matrix and related files to deconvolute bulk gene expression measures are available through Github (https://github.com/bakulskilab). Raw placental single-cell RNA-sequencing and raw placental bulk RNA-sequencing generated by this study are available in the Gene Expression Omnibus repository (accession number GSE182381). The placental single-cell RNA-sequencing data that support the findings of this study are available in the Database of Genotypes and Phenotypes (accession number phs001886.v1.p1) [16]. The preeclampsia case-control microarray data that support the findings of this study are available in Gene Expression Omnibus repository (accession number GSE75010) [32].

## Code Availability

All scripts to perform preprocessing, analyses, and deconvolution is available (https://github.com/bakulskilab).

