## Supplemental Material for "Placental gene expression-based cell type deconvolution: Cell proportions drive preeclampsia gene expression differences"

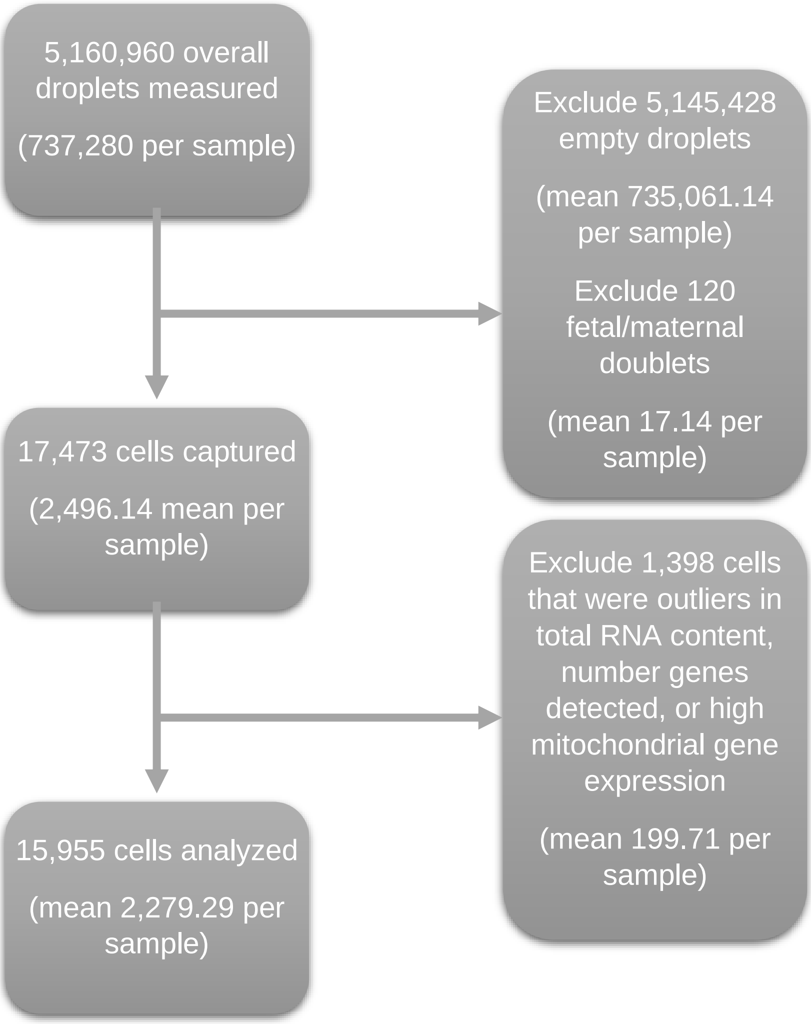


**Supplementary Figure 1.** Placental single-cell RNA sequencing quality control pipeline.

**Supplementary Table 1.** Summary of single-cell RNA-sequencing sample characteristics and sequencing quality metrics. Samples 1A/B and 2A/B were processed in our laboratory. Samples 3, 4, and 5 were downloaded from Pique-Regi et al., 2019 [1].

| | | Pre-filtering Quality Control Metrics | | | | | Post-filtering Quality Control Metrics | | | | | | | |
|---|---|---|---|---|---|---|---|---|---|---|---|---|---|---|
| Sample | Fetal Sex | Droplets Sequenced | Total Unique RNA Molecules | Total Unique Genes Detected | Total Cells | Maternal-Fetal Doublets Removed | Unique RNA Molecules (Median) | Unique RNA Molecules (IQR) | Unique Genes (Median) | Unique Genes (IQR) | Percent Mitochondria Gene Expression (Median) | Percent Mitochondria Gene Expression (IQR) | Putative Doublets Removed | Cells in Final Analytic Sample |
| 1A | F | 737,280 | 15,329,288 | 32,738 | 2,573 | 28 | 4,021 | 2,717 | 1,247 | 442 | 4.1 | 1.98 | 77 | 2,243 |
| 1B | F | 737,280 | 14,777,010 | 32,738 | 2,600 | 33 | 3,870 | 2,521 | 1,189 | 426 | 4.06 | 1.85 | 81 | 2,327 |
| 2A | M | 737,280 | 14,306,604 | 32,738 | 2,544 | 25 | 3,988 | 2,448 | 1,171 | 430 | 2.86 | 1.48 | 88 | 2,278 |
| 2B | M | 737,280 | 14,799,594 | 32,738 | 2,740 | 29 | 3,875 | 2,410 | 1,157 | 427 | 2.85 | 1.56 | 78 | 2,470 |
| 3 | M | 737,280 | 17,075,126 | 36,601 | 1,907 | 0 | 3,556 | 6,830 | 1,292 | 1,781 | 2.76 | 2.89 | 36 | 1,705 |
| 4 | F | 737,280 | 28,250,436 | 36,601 | 2,653 | 4 | 5,352 | 12,664 | 1,833 | 2,869 | 3.73 | 3.79 | 30 | 2,181 |
| 5 | M | 737,280 | 29,693,207 | 36,601 | 2,456 | 1 | 6,544 | 10,889 | 2,186 | 2,569 | 2.25 | 1.94 | 33 | 2,328 |

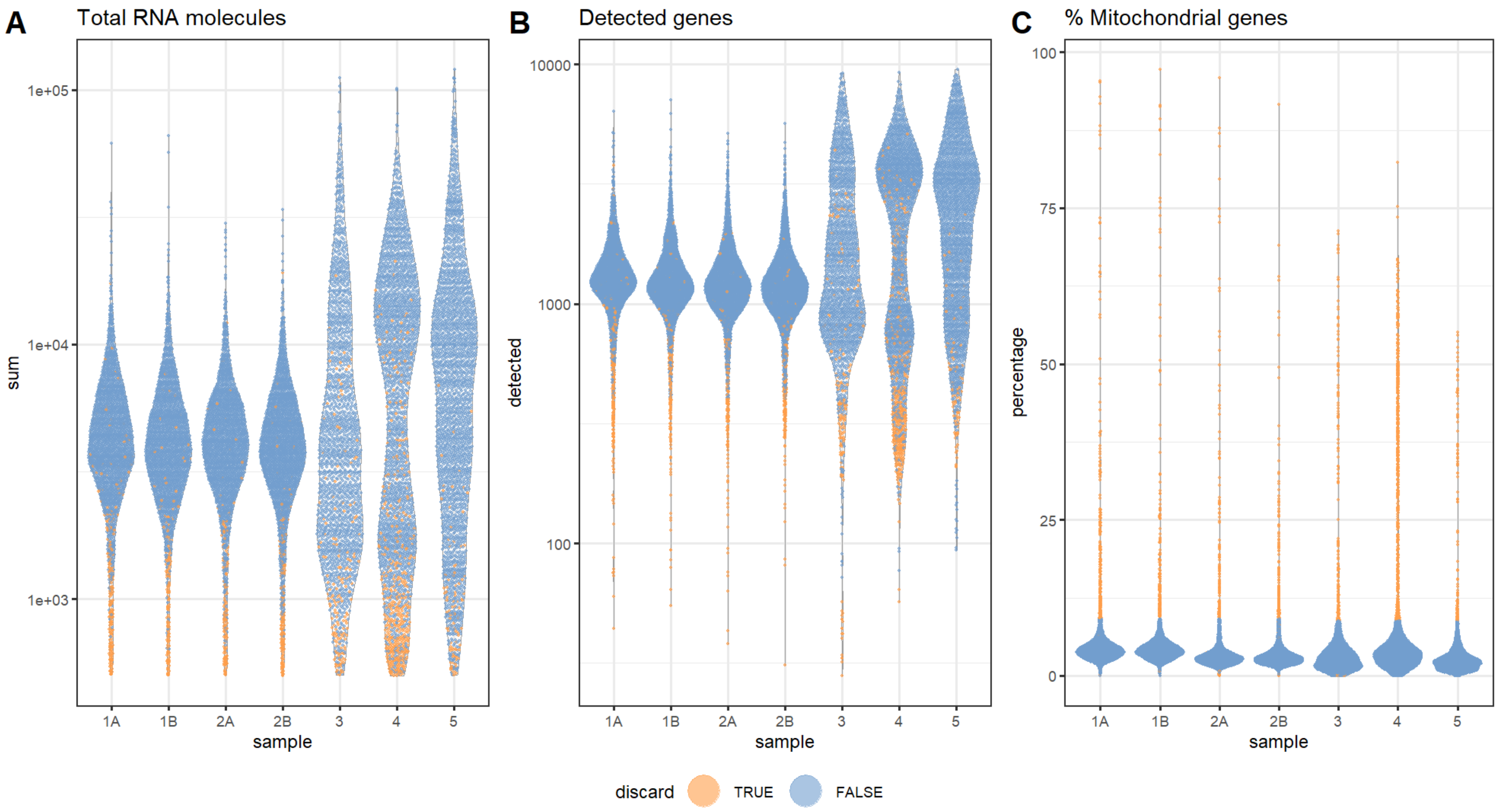


**Supplementary Figure 2.** Placental single cell RNA sequencing of quality metrics by sample, visualized using violin plots. Orange cells were discarded based on outlier status on any of the following metrics: (A) total RNA molecule count, (B) number of genes expressed, and (C) percent mitochondrial genes expressed.

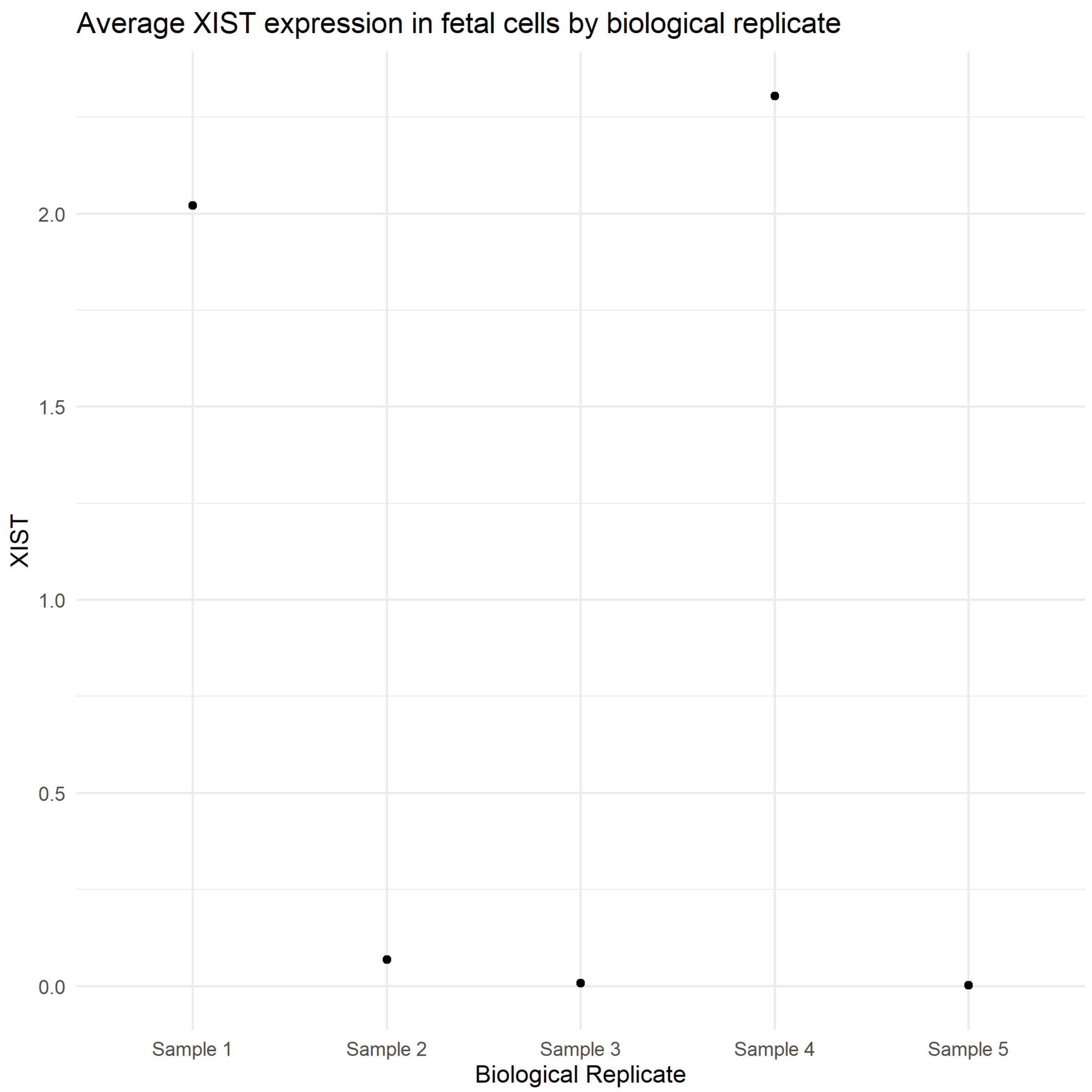


**Supplementary Figure 3.** Average *XIST* expression in fetal origin cells by biological replicates identifies Sample 1 as female due to high *XIST* expression, Sample 2 as male, and confirms fetal sex annotation for Samples 3, 4, and 5 as male, female, and male.

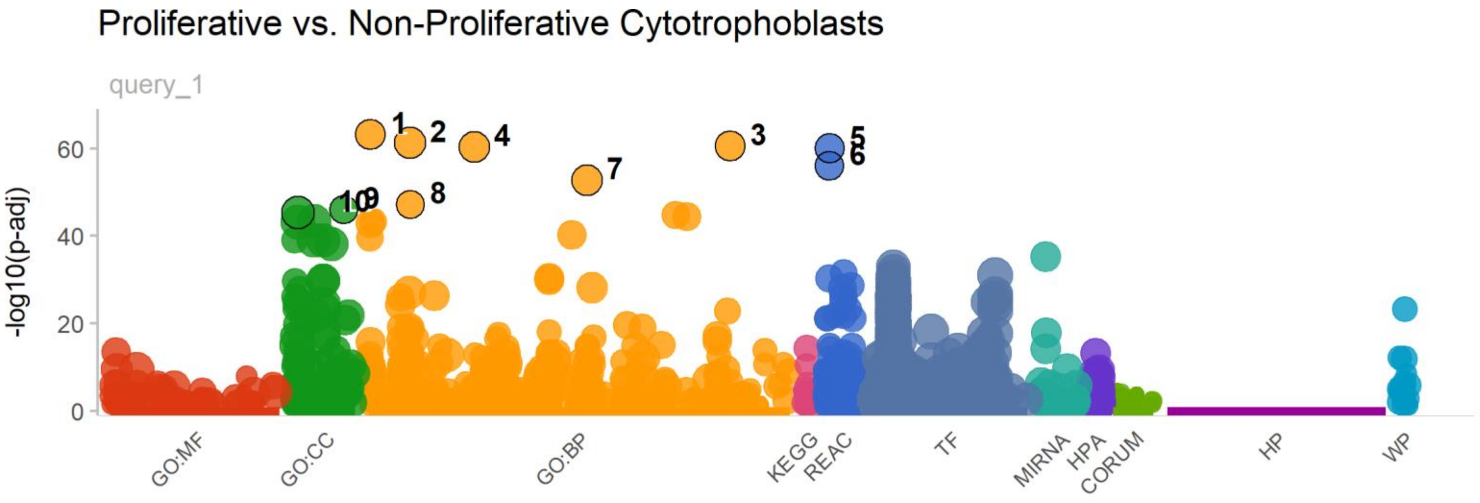


| id | source | term_id | term_name | term_size | | p_value |
|---|---|---|---|---|---|---|
| 1 | GO:BP | GO:0000278 | mitotic cell cycle | 957 | | 7.1e-64 |
| 2 | GO:BP | GO:0007049 | cell cycle | 1552 | | 5.4e-62 |
| 3 | GO:BP | GO:1903047 | mitotic cell cycle process | 829 | | 2.8e-61 |
| 4 | GO:BP | GO:0022402 | cell cycle process | 1307 | | 4.1e-61 |
| 5 | REAC | REAC:R-HSA-1640170 | Cell Cycle | 642 | | 9.1e-61 |
| 6 | REAC | REAC:R-HSA-69278 | Cell Cycle, Mitotic | 513 | | 9.1e-57 |
| 7 | GO:BP | GO:0051276 | chromosome organization | 1045 | | 1.5e-53 |
| 8 | GO:BP | GO:0007059 | chromosome segregation | 290 | | 5.3e-48 |
| 9 | GO:CC | GO:0098687 | chromosomal region | 280 | | 8.6e-47 |
| 10 | GO:CC | GO:0005654 | nucleoplasm | 4037 | | 4.3e-46 |

*g:Profiler (biit.cs.ut.ee/gprofiler)*

**Supplementary Figure 4.** Top 10 gene set enrichment results for proliferative vs. non-proliferative cytotrophoblasts differential expression results.

**Supplementary Table 2.** Top 10 marker gene expression (Bonferroni-adjusted p-value < .05) with highest average log fold change for each cell type cluster using Seurat's *FindAllMarkers* function. Table columns describe cluster cell type identity, marker gene symbol, average log fold change between this cell type cluster and other cell type clusters, prevalence of marker gene expression in cell type cluster, prevalence of marker gene expression in other cell type clusters, nominal p-value, and Bonferroni-adjusted p-value.

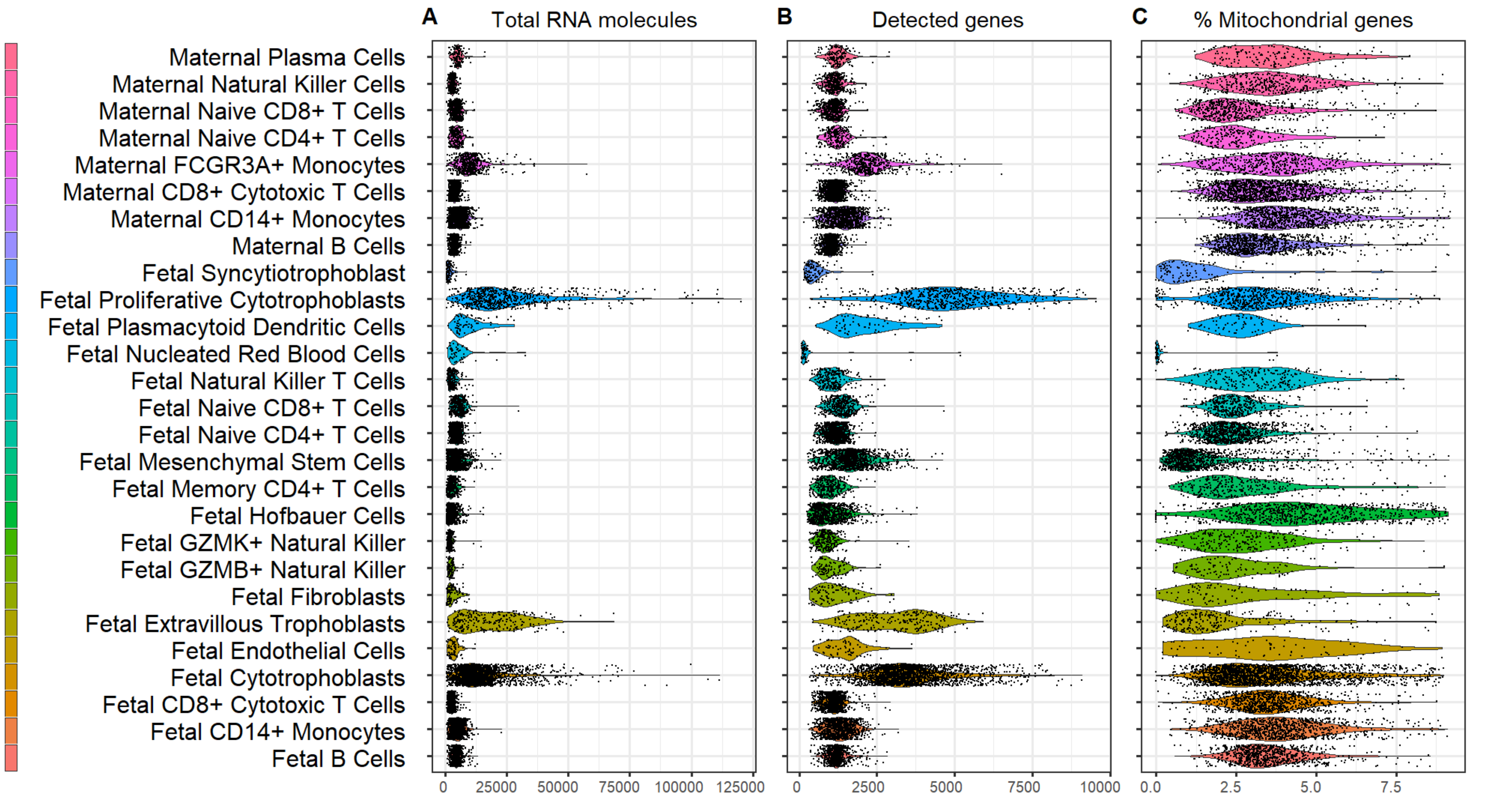


**Supplementary Figure 5.** Placental single cell RNA sequencing of quality metrics by cluster, visualized using violin plots. (A) Number of genes expressed, (B) total RNA molecule count, and (C) percent mitochondrial genes expressed.

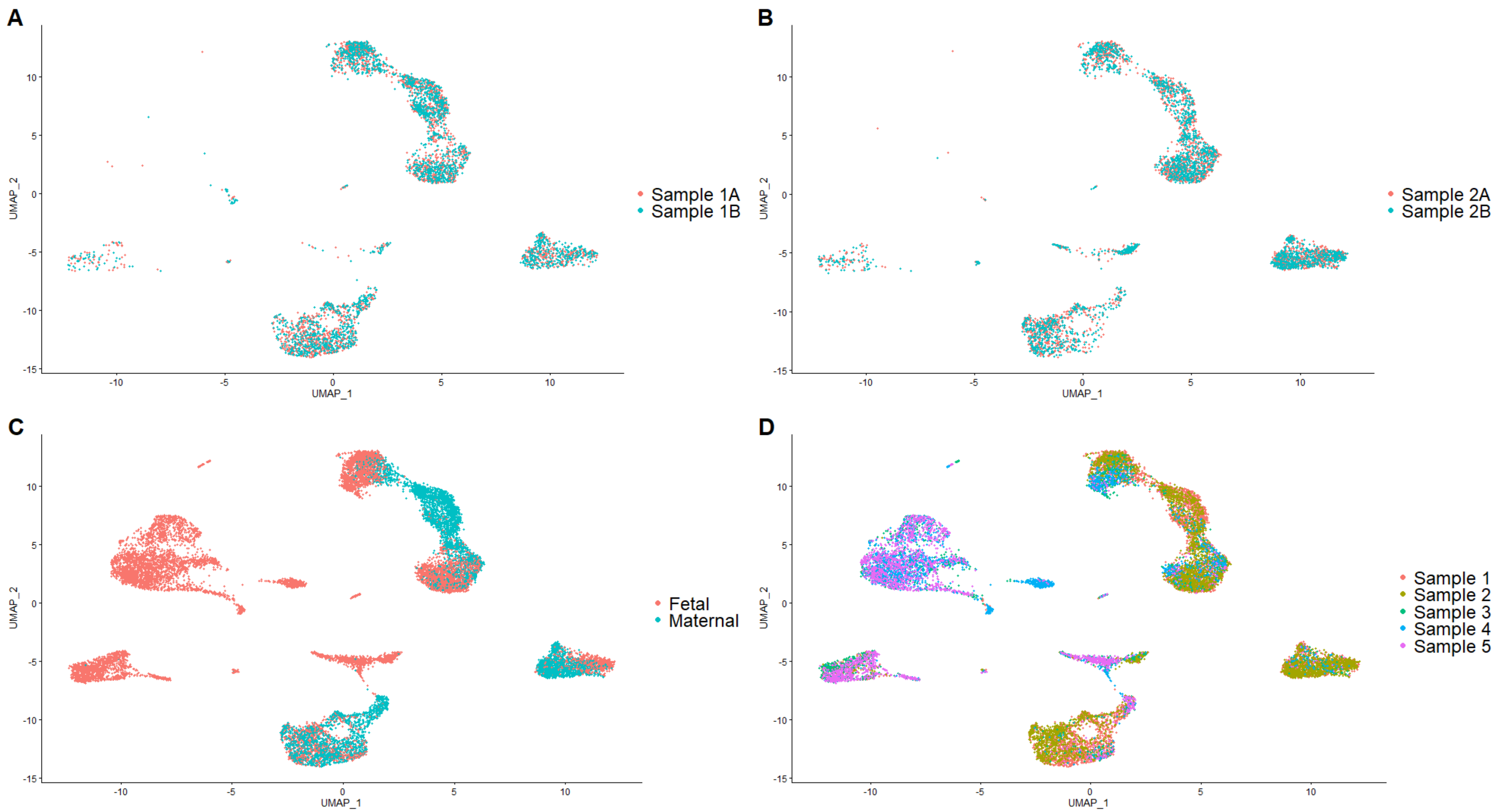


**Supplementary Figure 6.** Uniform Manifold Approximation and Projection (UMAP) plots colored by key variables. (A) Technical replication in Sample 1 with points colored by technical replicate. (B) Technical replication in Sample 2 with points colored by technical replicate. (C) Fetal/Maternal origin assignment by point color. (D) Biological replicates identified by point color with collapsed technical replicates.

**Supplementary Table 3.** Top ten results from g:Profiler2's g:GOSt functional enrichment function using cell type overexpressed genes from **Supplementary Table 2**. Table columns describe cell type cluster, gene ontology: biological process term name, false discovery rate-controlled (0.05) q-value, the size of the gene ontology term, and the number of top ranked genes used to test for enrichment against that term.

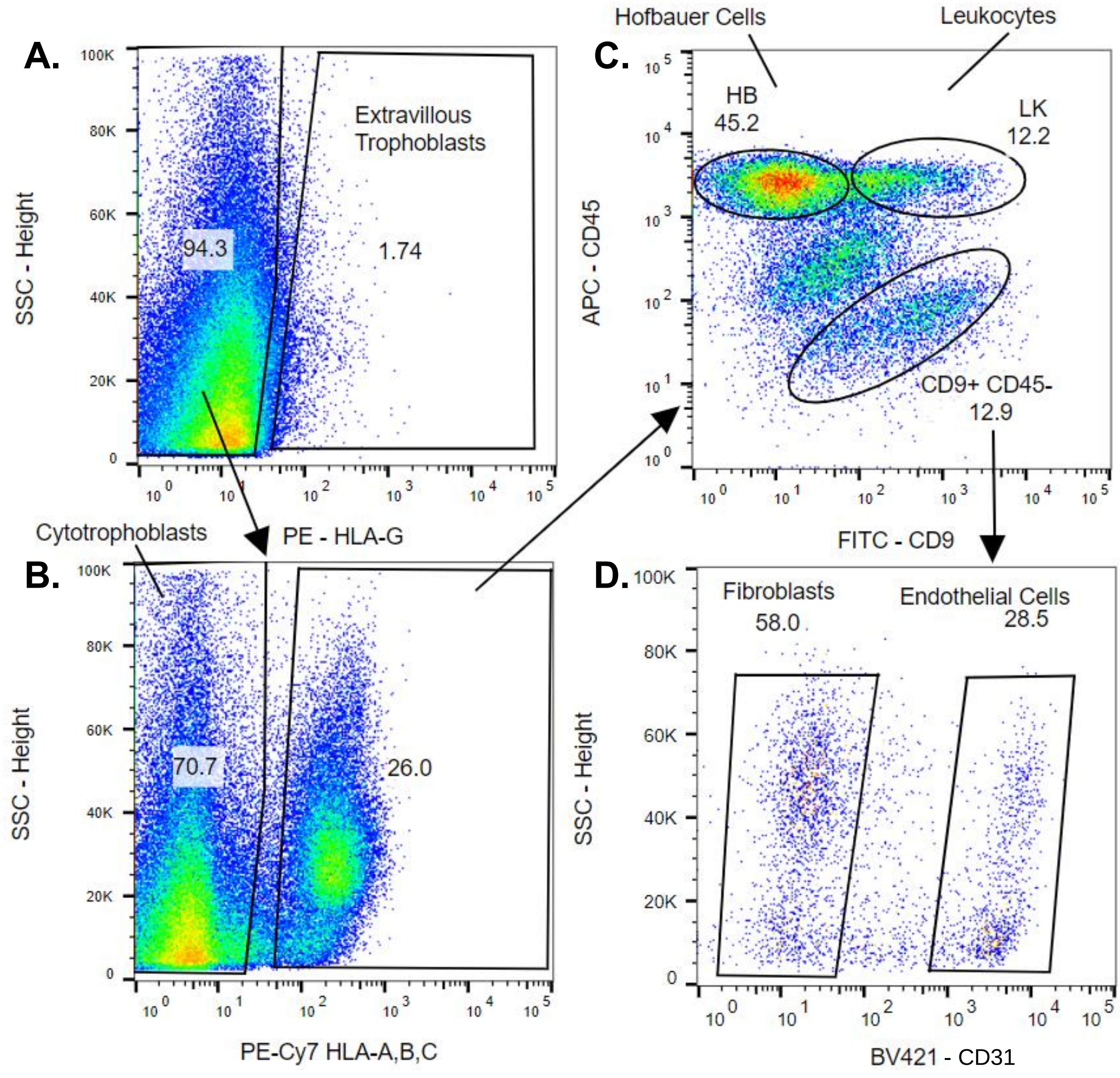


**Supplementary Figure 7.** A representative FACS sort. Gating strategy: (A) HLA-G/PE to positively select extravillous trophoblasts; (B) HLA-ABC/PE-CY7 to negatively enrich for cytotrophoblasts; (C) CD9/FITC by CD45/APC to positively select for Hofbauer cells and leukocytes; (D) CD31/BV421 to distinguish endothelial cells from CD31- fibroblasts.

**Supplementary Table 4.** Fluorescence-activated cell sorting and RNA-sequencing quality control results. Each sample was sorted into six cell type populations with a matched villous tissue sample. Cell count describes the total number of FACS-sorted cells. Table columns describe total RNA given in nanograms with RNA integrity index score (RIN), fastQC pass/fail, and whether the sample was sequenced in the paired-end (PE) or single-end (SE) and included in the experiment based on RIN score and total RNA (all sequenced samples included).
*Matched with sample 1 from the single-cell RNA-sequencing assay.

| Sample ID | Cell Type | Cell Count | Total RNA (ng) | RIN | fastQC | Sequenced (PE/SE) |
|---|---|---|---|---|---|---|
| 1* | Syncytiotrophoblast | N/A | 0.2 | 1 | N/A | Dropped |
| | Hofbauer | 1.57E+05 | 6.9 | 7.3 | Pass | SE |
| | Leukocyte | 2.00E+05 | 4.2 | 5 | Pass | SE |
| | Extravillous Trophoblast | 8.35E+03 | 1.1 | 6.4 | Pass | SE |
| | Cytotrophoblast | 2.83E+05 | 1.8 | 8.1 | Pass | SE |
| | Fibroblast | 1.55E+04 | 34 | 7.3 | Pass | SE |
| | Endothelial Cells | 7.60E+03 | 3 | 2.8 | N/A | Dropped |
| | Composite Tissue | N/A | 108 | 4.9 | Pass | SE |
| | Composite Tissue | N/A | 209 | 7.8 | Pass | PE |
| 6 | Syncytiotrophoblast | N/A | 0.2 | 1 | N/A | Dropped |
| | Hofbauer | 1.37E+05 | 63 | 8.3 | Pass | SE |
| | Leukocyte | 1.57E+05 | 1.6 | 4.9 | Pass | SE |
| | Extravillous Trophoblast | 3.60E+03 | 1.1 | 6 | Pass | SE |
| | Cytotrophoblast | 8.00E+04 | <0.1 | <0.1 | N/A | Dropped |
| | Fibroblast | 1.06E+04 | 35 | 8.1 | Pass | SE |
| | Endothelial Cells | 2.30E+03 | 2 | 8.1 | Pass | SE |
| | Composite Tissue | N/A | 332 | 8.7 | Pass | SE |
| 7 | Syncytiotrophoblast | N/A | 255 | 4 | Pass | SE |
| | Hofbauer | 1.02E+05 | 45 | 9.1 | Pass | SE |
| | Leukocyte | 8.30E+04 | 50 | 8.5 | Pass | SE |
| | Extravillous Trophoblast | 2.50E+04 | 7.2 | 2.8 | Pass | SE |
| | Cytotrophoblast | 3.72E+05 | 3.7 | 2.4 | N/A | Dropped |
| | Fibroblast | 3.61E+03 | 2.9 | 1.3 | N/A | Dropped |
| | Endothelial Cells | 2.97E+03 | 3 | 2.5 | N/A | Dropped |
| | Composite Tissue | N/A | 9572 | 5.3 | Pass | SE |
| 8 | Syncytiotrophoblast | N/A | 107 | 6.5 | Pass | PE |
| | Hofbauer | 8.40E+04 | 87 | 6.7 | Pass | PE |
| | Leukocyte | 2.20E+04 | 0.82 | 7.6 | Pass | PE |
| | Extravillous Trophoblast | 4.50E+04 | 2.4 | 8 | Pass | PE |
| | Cytotrophoblast | 2.56E+05 | 1.5 | 6.9 | Pass | PE |
| | Fibroblast | 1.70E+04 | 1.7 | 7.2 | Pass | PE |
| | Endothelial Cells | 1.20E+04 | 0.14 | 1 | Pass | Dropped |
| | Composite Tissue | N/A | 298 | 8.2 | Pass | PE |

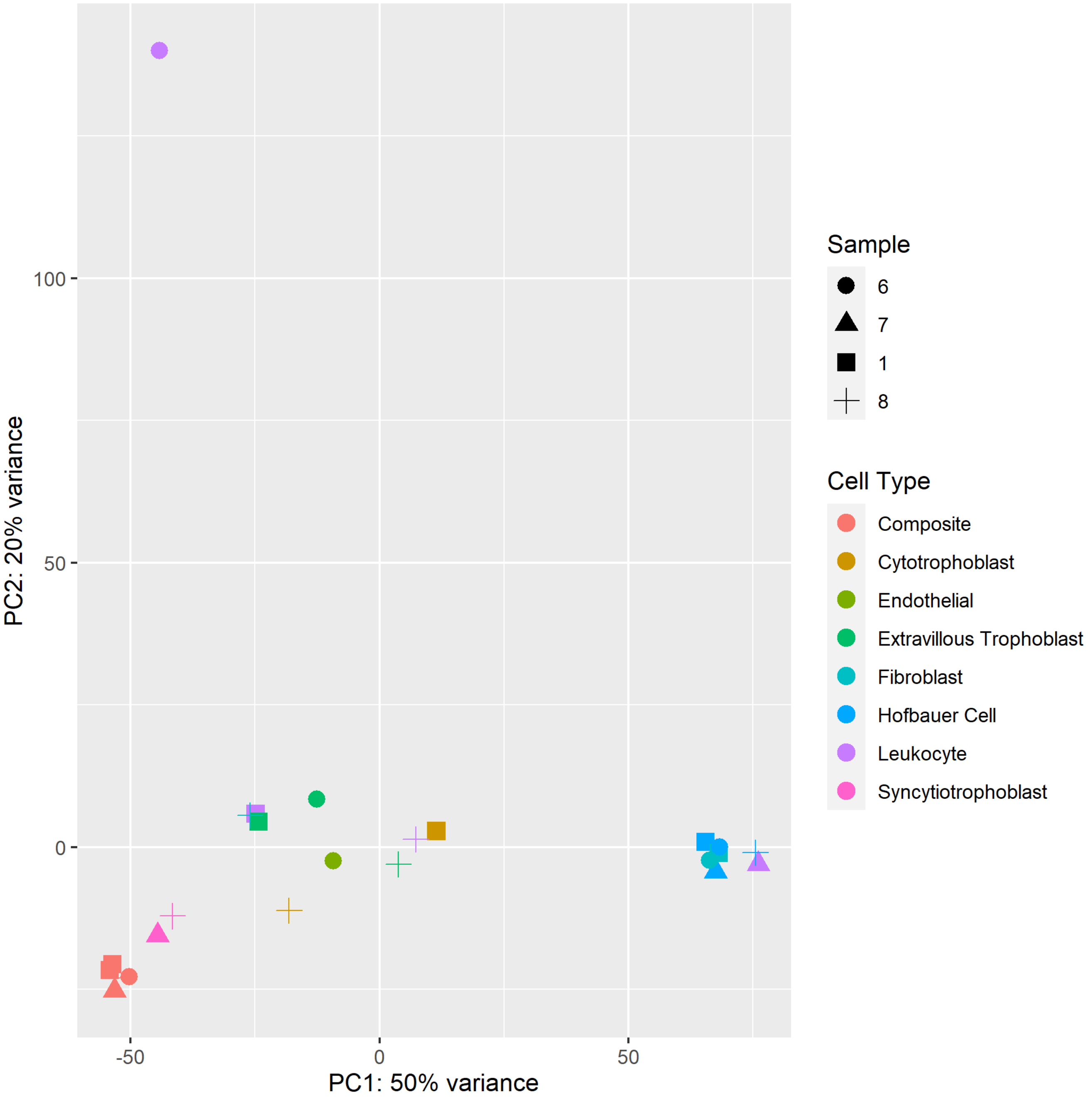


**Supplementary Figure 8.** Principal components plot of fluorescence-activated cell sorting bulk RNA-sequencing results on placentas 1, 6, 7, and 8. Point colors encode cell type. Composite represents whole villous tissue. Point shape denotes sample source.

**Supplementary Table 5.** Differentially upregulated genes sorted by descending test statistic from DESeq2 differential expression analysis comparing expression one placental cell type against average expression in other placental cell types, adjusted for sample source. Cell type describes the cell type of interest for the contrast. Gene refers to the name of the genomic feature testing. Log2 Fold-change is log2-transformed effect size of the tested gene. Base mean is the average normalized count values. P-value is the nominal p-value. q-value is the false discovery-controlled cutoff at 0.05.

**Supplementary Table 6.** Positive Gene Set Enrichment Analysis top ten results of differentially upregulated genes from DESeq2 differential expression analysis comparing one placental cell type against average expression in other placental cell types, adjusted for sample source. Cell Type: name of contrast, Name: name of Gene Ontology Biological Process gene set, Size: size of gene set, Normalized Enrichment Score: normalized gene set enrichment score, P-value: nominal p-value, False Discovery Rate-Controlled q-value: q-value with false discovery rate control at 0.05.

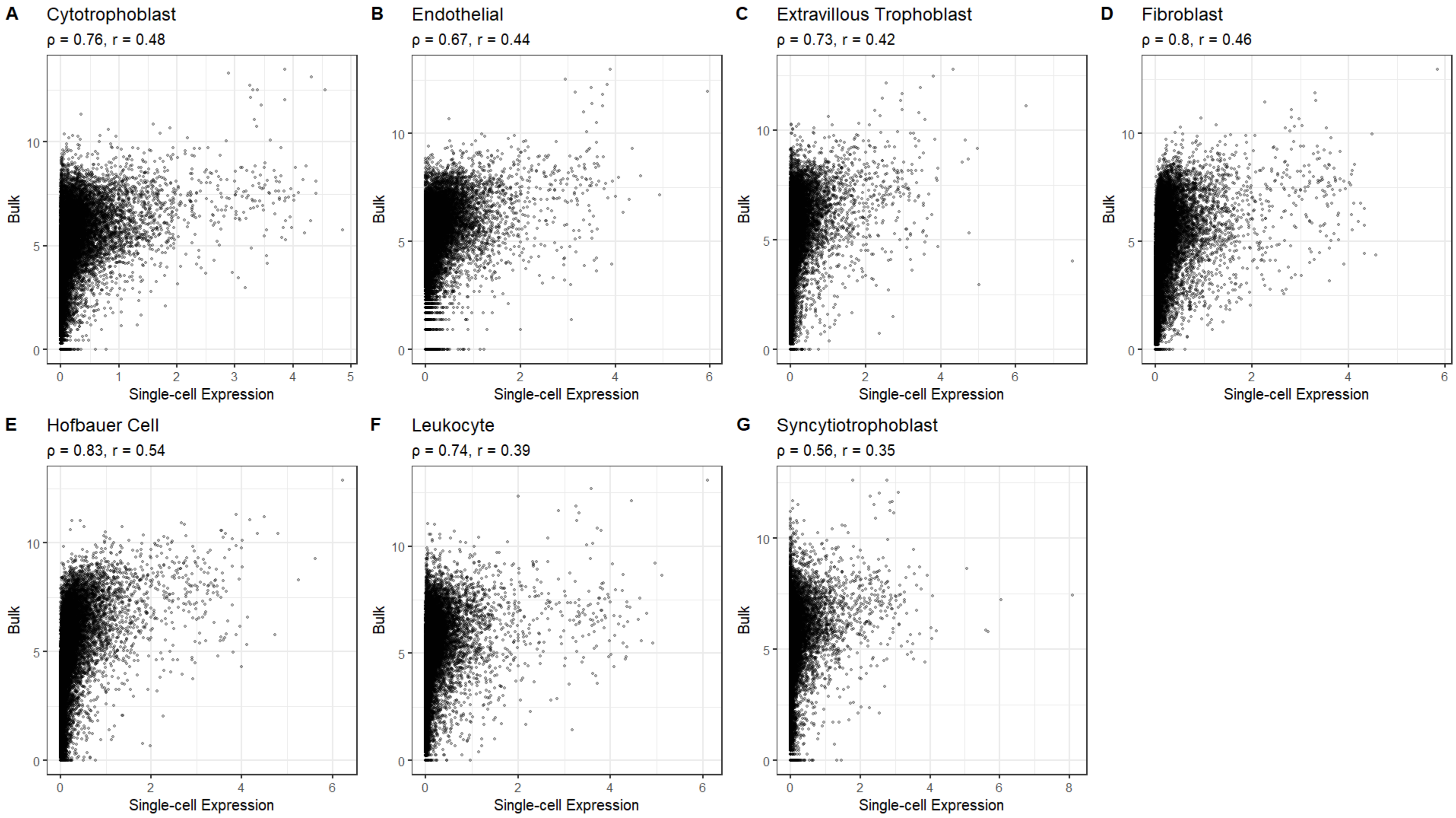


**Supplementary Figure 9.** Scatterplots comparing single-cell cluster and sorted cell type mean normalized gene expression across 27,490 common genes annotated with Spearman ($\rho$) and Pearson correlation (r) coefficients. X-axis encodes average normalized single-cell gene expression. Y-axis encodes average normalized bulk gene expression. (A) Cytotrophoblast. (B) Endothelial cells. (C) Extravillous trophoblasts. (D) Fibroblasts. (E) Hofbauer cells. (F) Leukocytes. (G) Syncytiotrophoblasts.

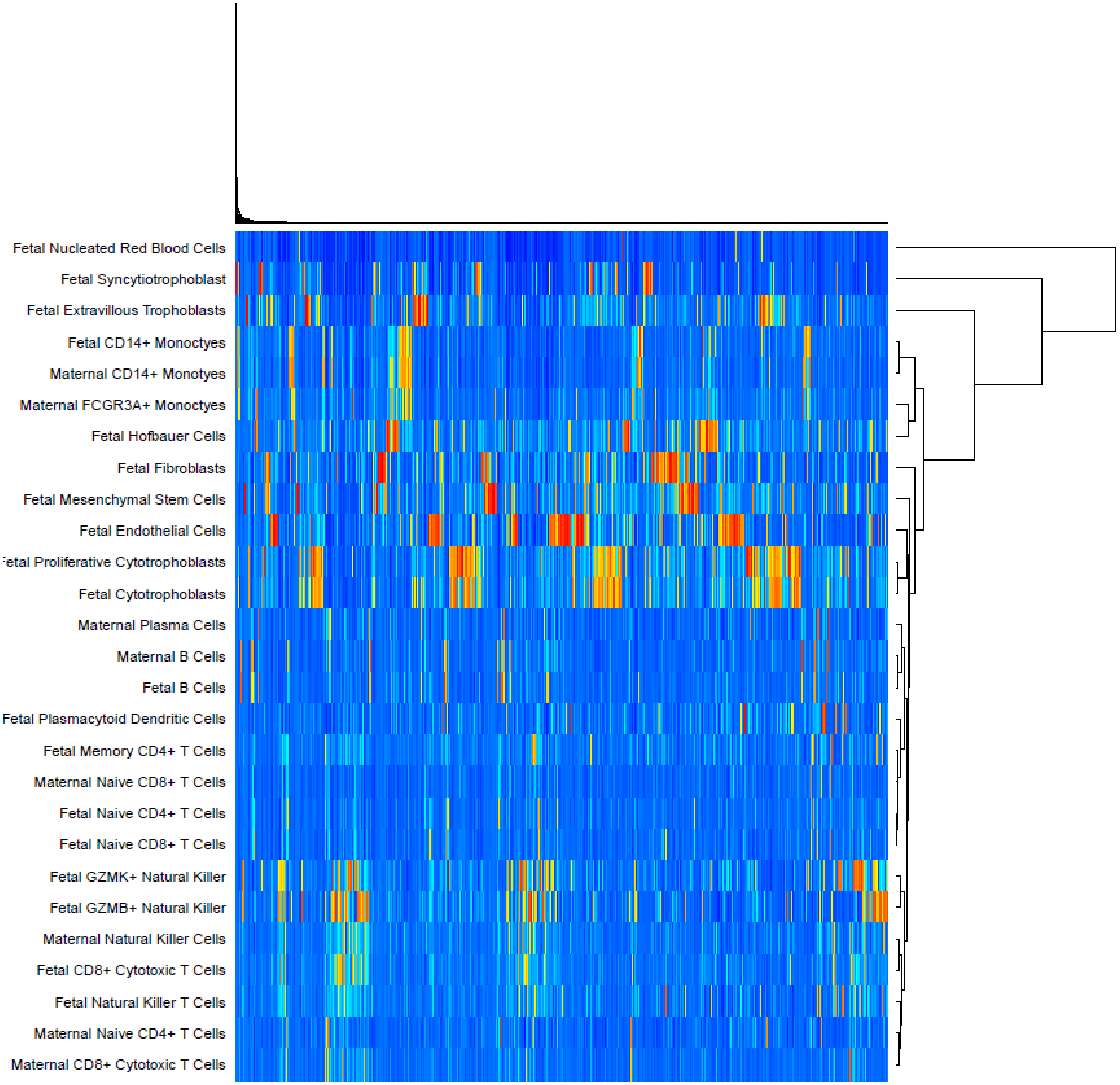


**Supplementary Figure 10**. Hierarchically clustered heatmap of signature gene expression matrix generated and used to deconvolute bulk placental tissue dataset. Cell types are encoded on the y-axis and genes are located along the x-axis. Blue indicates low expression of a gene and red represents high expression.

**Supplementary Table 7.** Estimated cell type proportions for preeclampsia bulk tissue dataset (GSE75010). Mixture corresponds to the individual microarray observations; each cell type is listed; p-value corresponds to the goodness-of-fit test for deconvolution (a statistically significant result suggests the signature gene reference matrix and the bulk mixture came from the same tissue); correlation describes the correlation between the original mixture and the estimated mixture among the signature genes; and RMSE is the root mean-square error between the original mixture and the imputed mixture among signature genes.

-

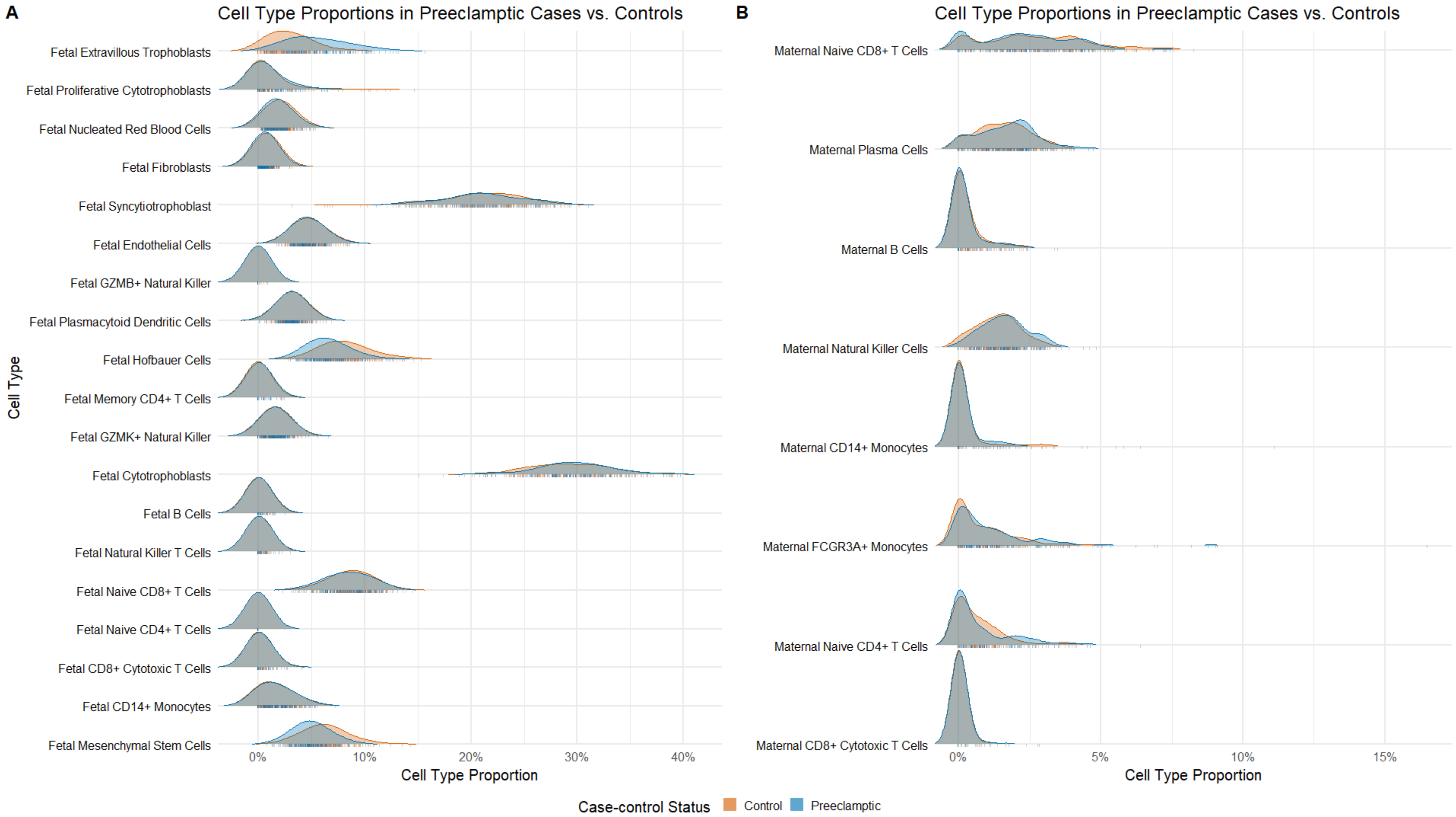


**Supplementary Figure 11.** Distribution of estimated cell type proportions in preeclamptic cases versus controls. Density distribution colored by case-control status. (A) Fetal cell types. (B) Maternal cell types.

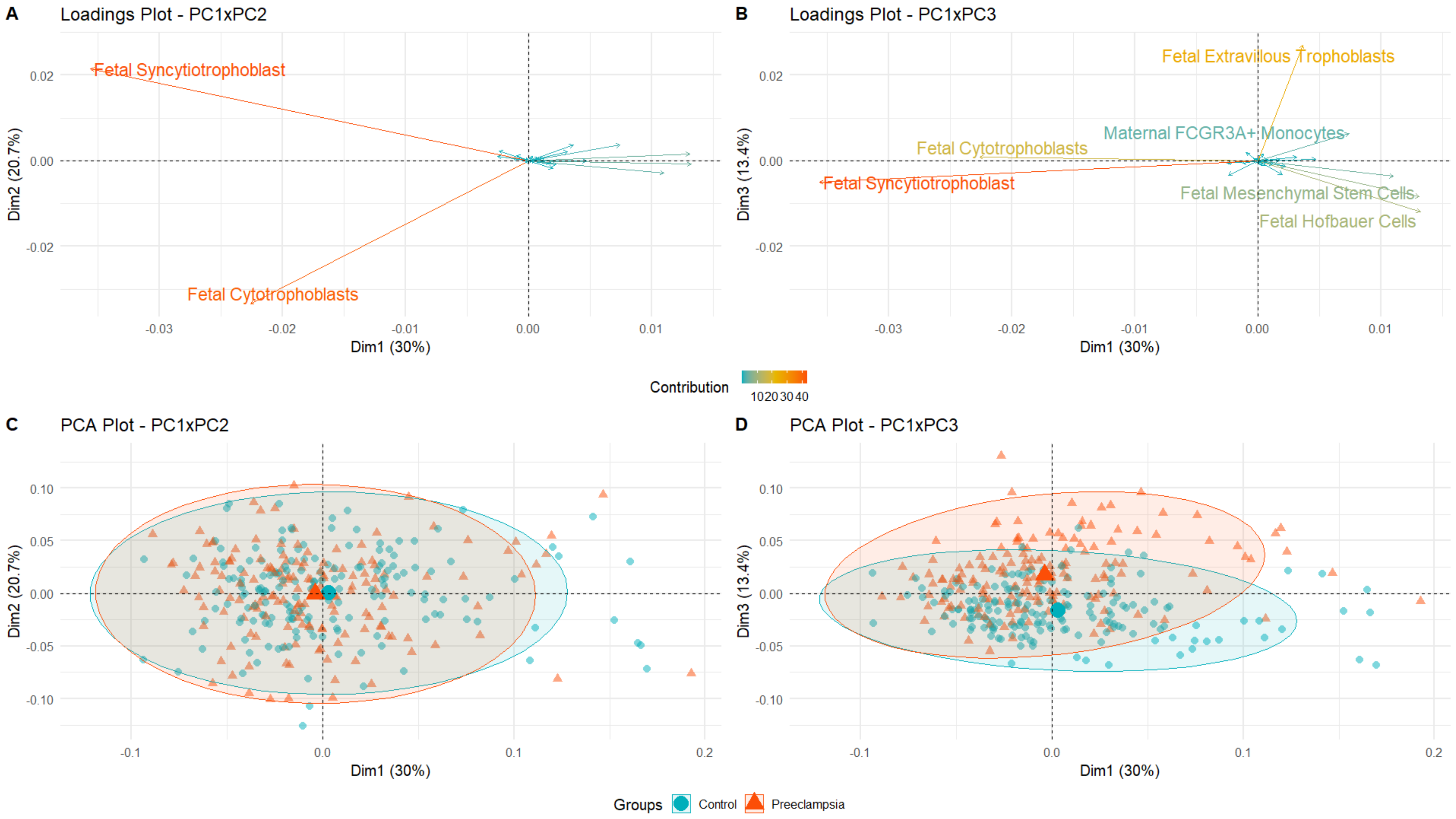


**Supplementary Figure 12.** Principal component results of estimated cell type proportions. (A) PC1 and PC2 dimension loadings dominated by fetal syncytiotrophoblasts and fetal cytotrophoblasts. (B) PC1 and PC3 dimension loadings. Fetal extravillous trophoblasts proportions are strongly correlated with PC3 whereas mesenchymal cell types fetal mesenchymal stem cells and fetal Hofbauer cells are correlated with PC1 and PC3. (C) Individual observations projected onto PC1xPC2 show little separation by preeclampsia case-control status. (D) Individual observations projected onto PC1xPC3 show moderate clustering by preeclampsia case-control status, particularly differentiated by PC3.
